# Preexisting heterogeneities in gene dosage sensitivity shaped sex chromosome evolution in mammals and birds

**DOI:** 10.1101/119016

**Authors:** Sahin Naqvi, Daniel W. Bellott, David C. Page

## Abstract

Mammalian X and Y chromosomes evolved from an ordinary autosomal pair; genetic decay decimated the Y, which in turn necessitated X chromosome inactivation (XCI). Genes of the ancestral autosomes are often assumed to have undertaken these transitions on uniform terms, but we hypothesized that they varied in their dosage constraints. We inferred such constraints from conservation of microRNA (miRNA)-mediated repression, validated by analysis of experimental data. X-linked genes with a surviving Y homolog have the most conserved miRNA target sites, followed by genes with no Y homolog and subject to XCI, and then genes with no Y homolog but escaping XCI; this heterogeneity existed on the ancestral autosomes. Similar results for avian Z-linked genes, with or without a W homolog, lead to a model of XY/ZW evolution incorporating preexisting dosage sensitivities of individual genes in determining their evolutionary fates, and ultimately shaping the mammalian and avian sex chromosomes.

## INTRODUCTION

The mammalian X and Y chromosomes evolved from a pair of ordinary autosomes over the past 300 million years (Lahn & Page, 1999). Only 3% of genes on the ancestral pair of autosomes survive on the human Y chromosome (Bellott et al., 2010; Skaletsky et al., 2003), compared to 98% on the X chromosome (Mueller et al., 2013). In females, one copy of the X chromosome is silenced by X inactivation (XCI); this silencing evolved on a gene-by-gene basis following Y gene loss (Jegalian & Page, 1998; Ross et al., 2005), and some genes escape XCI in humans (Carrel & Willard, 2005) and other mammals (Yang, Babak, Shendure, & Disteche, 2010). In parallel, the avian Z and W sex chromosomes evolved from a different pair of autosomes than the mammalian X and Y chromosomes (Bellott et al., 2010; Nanda et al., 1999; Ross et al., 2005).

Decay of the female-specific W chromosome was similarly extensive, but birds did not evolve a large-scale inactivation of Z-linked genes analogous to XCI in mammals (Itoh et al., 2007; Mank & Ellegren, 2009; Uebbing et al., 2015; Wright, Zimmer, Harrison, & Mank, 2015). Thus, genes previously found on the ancestral autosomes that gave rise to the mammalian or avian sex chromosomes have undergone significant changes in gene dosage. In modern mammals, these molecular events have resulted in three classes of ancestral X-linked genes representing distinct evolutionary fates: those with a surviving Y homolog, those with no Y homolog and subject to XCI, and those with no Y homolog but escaping XCI. In birds, two classes of ancestral Z-linked genes have arisen: those with or without a W homolog.

These classes of ancestral X- or Z-linked genes are not differentiated in existing models of sex chromosome evolution. Specifically, mathematically framed models of sex chromosome evolution do not consider the possibility that ancestral genes may differ with respect to their likelihood of surviving on the sex-specific Y or W chromosomes, or of acquiring dosage compensation following Y or W gene decay (Mullon, Wright, Reuter, Pomiankowski, & Mank, 2015). However, emerging evidence points towards critical heterogeneities in the properties of ancestral genes. For example, the trajectory of Y gene loss is consistent with exponential decay to a baseline level (Hughes et al., 2012), suggesting that mammalian Y chromosomes preferentially retained a subset of ancestral genes. Indeed, X- and Z-linked genes with surviving homologs on the mammalian Y or avian W chromosomes are enriched for important regulatory functions and predictors of haploinsufficiency compared to those lacking Y or W homologs (Bellott et al., 2014, 2017). These observations led us to ask whether the three classes of X-linked genes and the two classes of Z-linked genes show molecular signatures of dosage sensitivity that can be traced back to their ancestral, autosomal precursors.

Robust analyses of the selective pressures underlying either gene loss from the sex-specific chromosomes or the subsequent acquisition of XCI require precise delineation of the set of genes that were present on the ancestral autosomes. We took advantage of our previous reconstructions of the ancestral gene content of both the mammalian and avian sex chromosomes achieved through sequencing and bioinformatics approaches (Bellott et al., 2014, 2017, 2010; Hughes et al., 2012; Mueller et al., 2013). We reasoned that if X-linked genes, or Z-linked genes, differ in their dosage sensitivities, they also may differ in their regulatory states, with stronger purifying selection acting on regulatory elements associated with dosage-sensitive genes. We focused on microRNAs (miRNAs), small noncoding RNAs that frequently function as “tuners” of gene dosage by lowering target mRNA levels through pairing to the 3’ untranslated region (UTR) (Bartel, 2009). miRNA targets can be predicted genome-wide and followed through evolutionary time, enabling robust estimates of site-specific conservation (Friedman, Farh, Burge, & Bartel, 2009) and allowing us to reconstruct the regulatory states of the autosomal ancestors of current X- and Z-linked genes.

First, through genome-wide analysis of human copy number variation, we refined and validated a method for inferring dosage sensitivity based on patterns of conserved miRNA targeting. Using this tool, we inferred heterogeneities in dosage sensitivity between the three classes of X-linked genes and the two classes of Z-linked genes. We then demonstrated that these differences were likely to have been present on the ancestral autosomes that gave rise to the mammalian and avian sex chromosomes. Finally, we used publically available experimental datasets to validate the efficacy, in living cells, of the conserved miRNA targeting. These findings have important implications for our understanding of X and Z dosage compensation, and provide the foundation for a model of X-Y and Z-W evolution in which both survival on the sex-specific Y or W chromosome and the subsequent evolution of XCI in mammals were determined by preexisting sensitivities to under- and overexpression.

## RESULTS

### Analysis of human copy number variation indicates conserved microRNA targeting of dosage-sensitive genes

We first sought to determine whether conserved targeting by microRNAs (miRNAs) correlates with dosage sensitivity across the human genome. To estimate pressure to maintain miRNA targeting, we used published probabilities of conserved targeting (P_CT_ scores) for each gene-miRNA interaction in the human genome. Since the PCT score intrinsically controls for differences in the background conservation and sequence composition of individual 3’UTRs, it is comparable across both genes and miRNA families (Friedman et al., 2009). We refer to these P_CT_ scores as “miRNA conservation scores” in the remainder of the text.

Genes for which increases in dosage are deleterious should be depleted from the set of observed gene duplications in healthy human individuals. We used a catalogue of rare genic copy number variation among 59,898 control human exomes (Exome Aggregation Consortium, ExAC)(Ruderfer et al., 2016) to classify autosomal protein-coding genes as exhibiting or lacking duplication or deletion in healthy individuals (see Methods). We compared duplicated and non-duplicated genes with the same deletion status in order to control for differences in sensitivity to underexpression. We found that non-duplicated genes have significantly higher miRNA conservation scores than duplicated genes, irrespective of deletion status (Figure 1A,B). Non-deleted genes also have significantly higher scores than deleted genes irrespective of duplication status (Figure 1 – figure supplement 1), but duplication status has a greater effect on miRNA conservation scores than does deletion status (Figure 1 – figure supplement 2). Thus, conserved miRNA targeting is a feature of genes sensitive to changes in gene dosage in humans and is especially informative with regards to sensitivity to overexpression.

**Figure 1:**
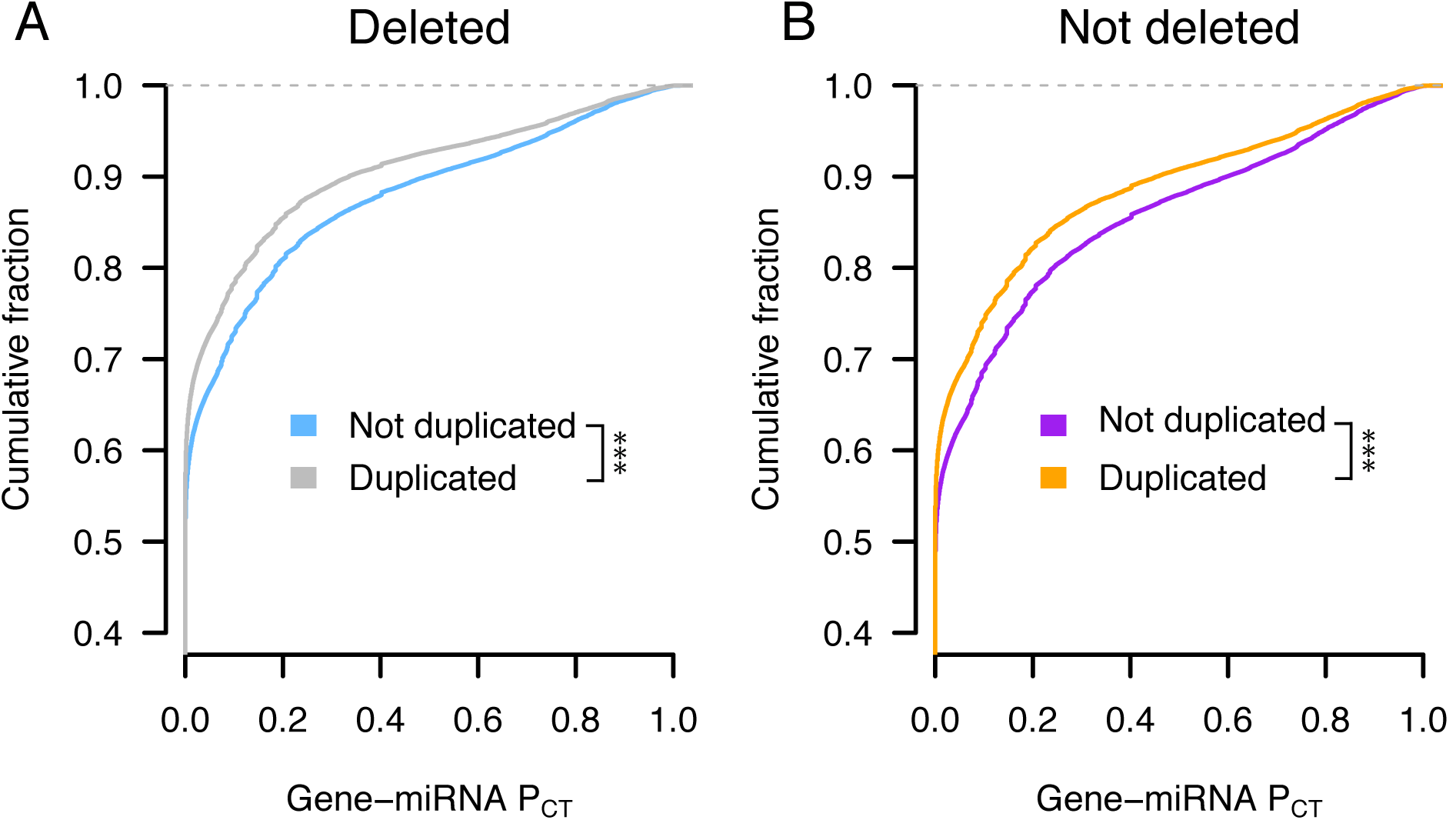
Conserved miRNA targeting of autosomal genes stratified by copy number variation in 59,898 human exomes. Probabilities of conserved targeting (P_CT_) of all gene-miRNA interactions involving non-duplicated and duplicated genes, further stratified as (A) deleted (blue, n = 80,290 interactions from 3,976 genes; grey, n = 69,339 interactions from 4,118 genes) or (B) not deleted (purple, n = 72,826 interactions from 3,510 genes; orange, n = 51,514 interactions from 2,916 genes). *** p < 0.001, two-sided Kolmogorov-Smirnov test.

### X-Y pairs and X-inactivated genes have higher miRNA conservation scores than X escape genes

We next assessed whether the three classes of X-linked genes differ with respect to dosage sensitivity as inferred by conserved miRNA targeting. To delineate these classes, we began with the set of ancestral genes reconstructed through cross-species comparisons of the human X chromosome and orthologous chicken autosomes (Bellott et al., 2014, 2017, 2010; Hughes et al., 2012; Mueller et al., 2013). We designated ancestral X-linked genes with a surviving human Y homolog (Skaletsky et al., 2003) as X-Y pairs and also considered the set of X-linked genes with a surviving Y homolog in any of eight mammals (Bellott et al., 2014) to increase the phylogenetic breadth of findings regarding X-Y pairs. A number of studies have catalogued the inactivation status of X-linked genes in various human tissues and cell-types. We used a meta-analysis that combined results from three studies by assigning a “consensus” X-inactivation status to each gene (Balaton, Cotton, & Brown, 2015) to designate the remainder of ancestral genes lacking a Y homolog as subject to or escaping XCI. In summary, we classified genes as either: 1) X-Y pairs, 2) lacking a Y homolog and subject to XCI (X-inactivated), or 3) lacking a Y homolog but escaping XCI (X escape).

We found that human X-Y pairs have the highest miRNA conservation scores, followed by X-inactivated and finally X escape genes (Figure 2A). The expanded set of X-Y pairs across eight mammals also has significantly higher miRNA conservation scores than ancestral X-linked genes with no Y homolog (Figure 2B). Observed differences between miRNA conservation scores are not driven by distinct subsets of genes in each class, as indicated by gene resampling with replacement (Figure 2 – figure supplement 1), and are not accounted for by differences in haploinsufficiency, as indicated by comparisons of gene subsets matched by published estimates of haploinsufficiency probabilities (Huang, Lee, Marcotte, & Hurles, 2010) (Figure 2 – figure supplement 2). Furthermore, the decrease in miRNA conservation scores of X escape genes relative to X-inactivated genes and X-Y pairs is not driven by genes that escape variably across individuals (Figure 2 –figure supplement 3). Together, these results indicate heterogeneity in sensitivity to both under- and overexpression among the three classes of ancestral X-linked genes: X-Y pairs are the most dosage-sensitive, while X-inactivated genes are of intermediate dosage sensitivity, and X escape genes are the least dosage-sensitive.

**Figure 2.**
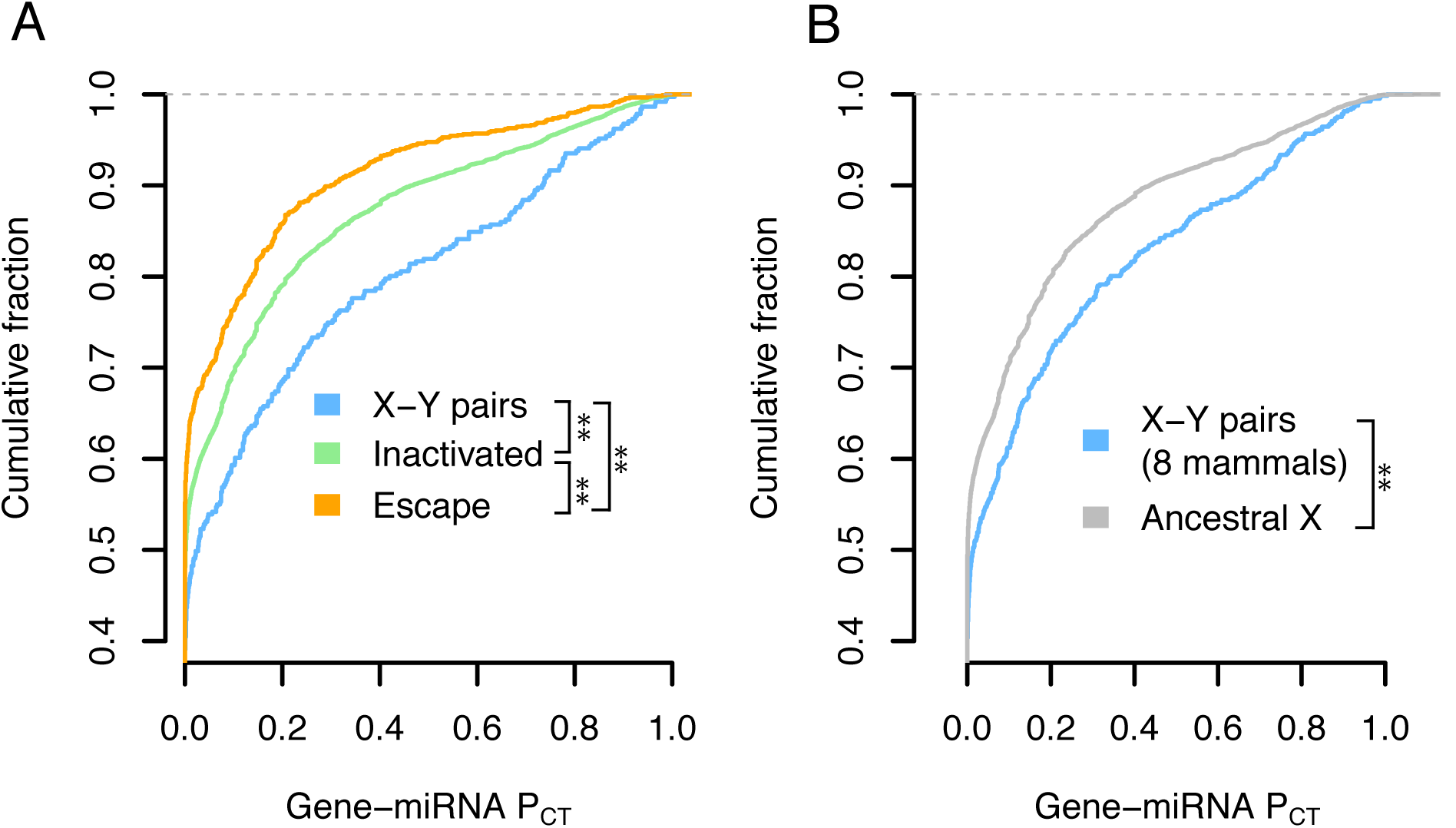
X-Y pairs and X-inactivated genes have higher miRNA conservation scores than X escape genes. a) P_CT_ score distributions of all gene-miRNA interactions involving (A) human X-Y pairs (n = 371 interactions from 15 genes), X-inactivated genes (n = 6,743 interactions from 329 genes), and X escape genes (n = 1,037 interactions from 56 genes), or (B) X-Y pairs across eight sequenced mammalian Y chromosomes (n = 647 interactions from 32 genes) and other ancestral X genes (n = 8,831 interactions from 457 genes). ** p < 0.01, two-sided Kolmogorov-Smirnov test.

### Patterns of X-linked miRNA targeting were present on the ancestral autosomes

We next asked whether the differences in miRNA targeting for the three classes of X-linked genes were present on the ancestral autosomes that gave rise to the mammalian X and Y chromosomes. We focused on miRNA target sites in the 3’ UTR of human orthologs that align with perfect identity to a site in the chicken genome; these sites were likely present in the common ancestor of mammals and birds and thus likely present on the ancestral autosomes. We found that X-Y pairs have the highest fraction of genes with at least one human-chicken-conserved target site, followed by X-inactivated genes, and then X escape genes, with similar results for the expanded set of X-Y pairs across 8 mammals (Figure 3A,B). We observed the same pattern after accounting for the background conservation of each individual 3’ UTR (see Methods) (Figure 3 – figure supplement 1). These results indicate that the autosomal precursors of X-Y pairs and X-inactivated genes were more dosage-sensitive than those of X escape genes; present-day heterogeneities in dosage sensitivity on the mammalian X chromosome were present on the ancestral autosomes from which it derived.

**Figure 3.**
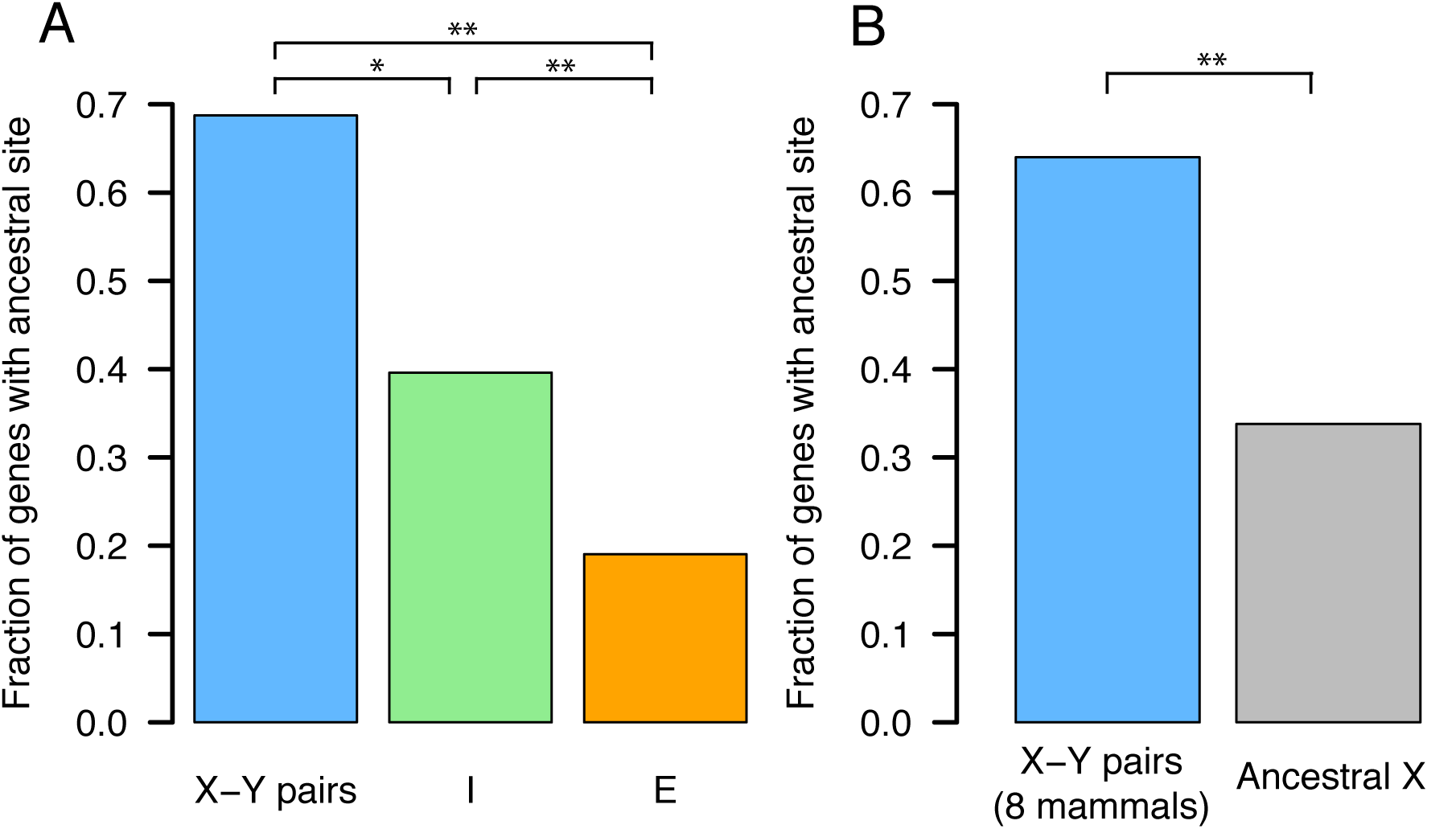
Patterns of X-linked miRNA targeting were present on the ancestral autosomes. Each bar represents the fraction of genes with at least one miRNA target site found in the 3’ UTR of both human and chicken orthologs. (A) Human X-Y pairs (n = 16), X-inactivated genes (n = 251), and X escape genes (n = 42). (B) X-Y pairs across eight sequenced mammalian Y chromosomes (n = 25) and other ancestral X genes (n = 351). * p < 0.05, ** p < 0.01, two-sided Fisher’s exact test.

### Z-W pairs have higher miRNA conservation scores than other ancestral Z-linked genes

We recently showed that avian Z-linked genes with surviving homologs on the female-specific W chromosome (Z-W pairs) are enriched for predictors of haploinsufficiency relative to those lacking a W homolog, indicating sensitivity to underexpression (Bellott et al., 2017). Our analyses of X-linked miRNA targeting presented here document heterogeneities in sensitivity to both under- and overexpression; we therefore assessed whether Z-W pairs are similarly sensitive to both increases and decreases in gene dosage. We used the set of ancestral genes reconstructed through cross-species comparisons of the avian Z chromosome and orthologous human autosomes and focused on the set of Z-W pairs identified by sequencing of the chicken W chromosome (Bellott et al., 2017, 2010). To increase the phylogenetic breadth of our comparisons, we also included candidate Z-W pairs obtained through comparisons of male and female genome assemblies (4 species set) or inferred by read-depth changes in female genome assemblies (14 species set, see Methods for details) (Zhou et al., 2014). Since the more complete 3’UTR annotations in the human genome relative to chicken allow for a more accurate assessment of conserved miRNA targeting, we analyzed the 3’ UTRs of the human orthologs of chicken Z-linked genes, reasoning that important properties of chicken Z-linked genes should be retained by their human autosomal orthologs.

We found that the human orthologs of Z-W pairs have higher miRNA conservation scores than the human orthologs of other ancestral Z genes (Figure 3A). Differences in miRNA conservation scores between Z-W pairs and other ancestral Z genes remained significant when considering the expanded sets of Z-W pairs across four and 14 avian species (Figure 3B,C). These differences are not driven by distinct subsets of genes (Figure 4 – figure supplement 1) and are not accounted for by haploinsufficiency probability (Figure 4 – figure supplement 2). We infer that Z-linked genes with a surviving W homolog are more sensitive to changes in dosage — both increases and decreases — than are genes without a surviving W homolog.

**Figure 4.**
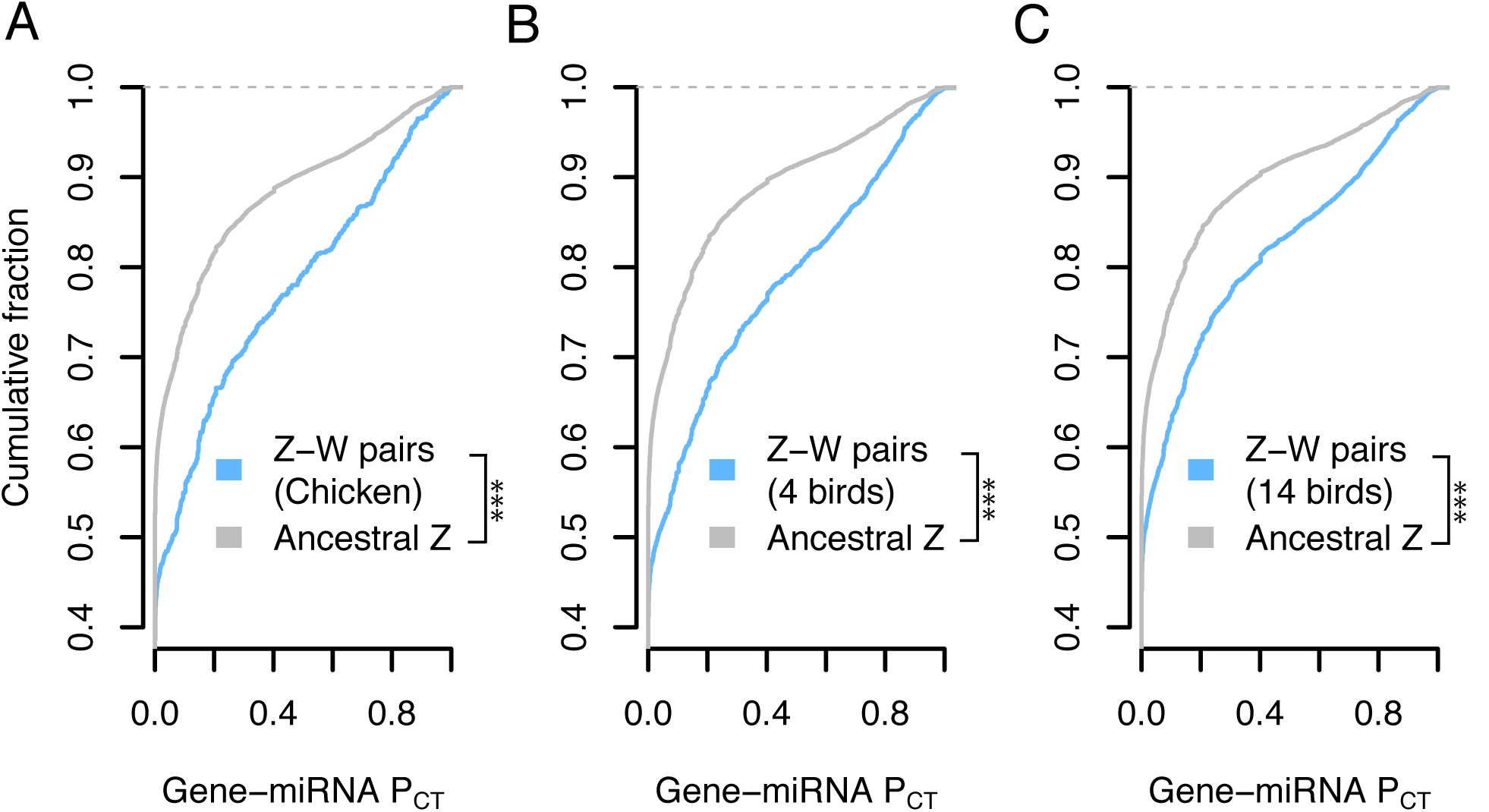
Z-W pairs have higher miRNA conservation scores than other ancestral Z-linked genes. P_CT_ score distributions of all gene-miRNA interactions involving the human orthologs of (A) chicken Z-W pairs (n = 832 interactions from 28 genes) and other ancestral Z genes (n = 16,692 interactions from 657 genes), (B) Z-W pairs including predictions from three additional birds with male and female genome sequence (n = 2,187 interactions from 78 genes) and other ancestral Z genes (n = 15,357 interactions from 607 genes), or (C) Z-W pairs including read depth-based predictions from 10 additional birds with only female genome sequence (n = 4,458 interactions from 157 genes) and other ancestral Z genes (n = 13,086 interactions from 528 genes). *** p < 0.001, two-sided Kolmogorov-Smirnov test.

### Patterns of Z-linked miRNA targeting were present on the ancestral autosomes

We next asked whether the differences in miRNA targeting between Z-W pairs and other ancestral Z-linked genes were present on the ancestral autosomes that gave rise to the avian Z and W chromosomes. We found that both chicken Z-W pairs and the predicted 4- and 14-species sets of Z-W pairs are enriched for human-chicken-conserved miRNA target sites relative to their Z-linked counterparts without surviving W homologs (Figure 5A–C), even when accounting for the background conservation of each individual 3’ UTR (Figure 5 – figure supplement 1). Thus, the autosomal precursors of avian Z-W pairs were more dosage-sensitive than the autosomal precursors of Z-linked genes that lack a W homolog.

**Figure 5.**
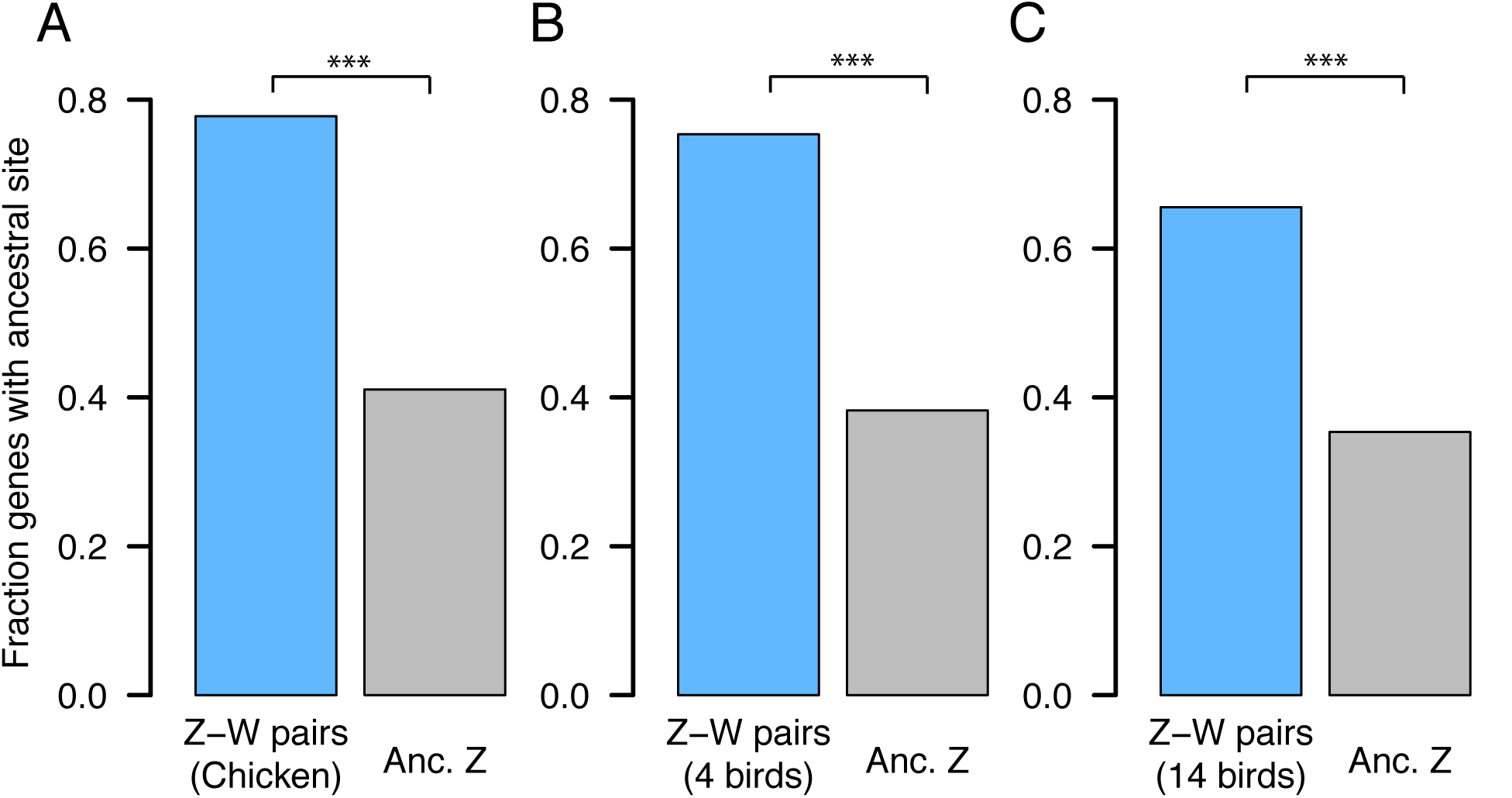
Patterns of Z-linked miRNA targeting were present on the ancestral autosomes. Each bar represents the fraction of genes with at least one miRNA target site found in the 3’ UTR of both human and chicken orthologs. (A) Chicken Z-W pairs (n = 27) and other ancestral Z genes (n = 578). (B) Z-W pairs including predictions from three additional birds with male and female genome sequence (n = 73) and other ancestral Z genes (n = 532). (C) Z-W pairs including read depth-based predictions from 10 additional birds with only female genome sequence (n = 147) and other ancestral Z genes (n = 458). * p < 0.05, ** p < 0.01, two-sided Fisher’s exact test.

### Analyses of experimental datasets validate miRNA target site efficacy

Our results to this point, which indicate heterogeneities in dosage constraints among X- or Z-linked genes as inferred by predicted miRNA target sites, lead to predictions regarding the efficacy of these sites in vivo. To test these predictions, we turned to publically available experimental datasets consisting both of gene expression profiling following transfection or knockout of individual miRNAs, and of high-throughput crosslinking-immunoprecipitation (CLIP) to identify sites that bind Argonaute in vivo (see Methods). If the above-studied sites are effective in mediating target repression, targets of an individual miRNA should show increased (decreased) expression levels or Argonaute binding following miRNA transfection (knockout). Together, our analyses of publically available datasets fulfilled these predictions, validating the efficacy of sites targeting ancestral X-linked genes and the autosomal orthologs of ancestral Z-linked genes in multiple cellular contexts and species (Figure 6). From the gene expression profiling data, we observed results consistent with effective targeting by a) ten different miRNA families in human HeLa cells (Figure 6 – figure supplement 1), b) four different miRNAs in human HCT116 and HEK293 cells (Figure 6 – figure supplement 2), and c) miR-155 in mouse B and Th1 cells (Figure 6 – figure supplement 3). In the CLIP data, the human orthologs of X- or Z-linked targets of miR-124 are enriched for Argonaute-bound clusters that appear following miR-124 transfection, while a similar but non-significant enrichment is observed for miR-7 (Figure 6 – figure supplement 4). Thus, conserved miRNA target sites used to infer dosage constraints on X-linked genes and the autosomal orthologs of Z-linked genes can effectively mediate target repression.

**Figure 6.**
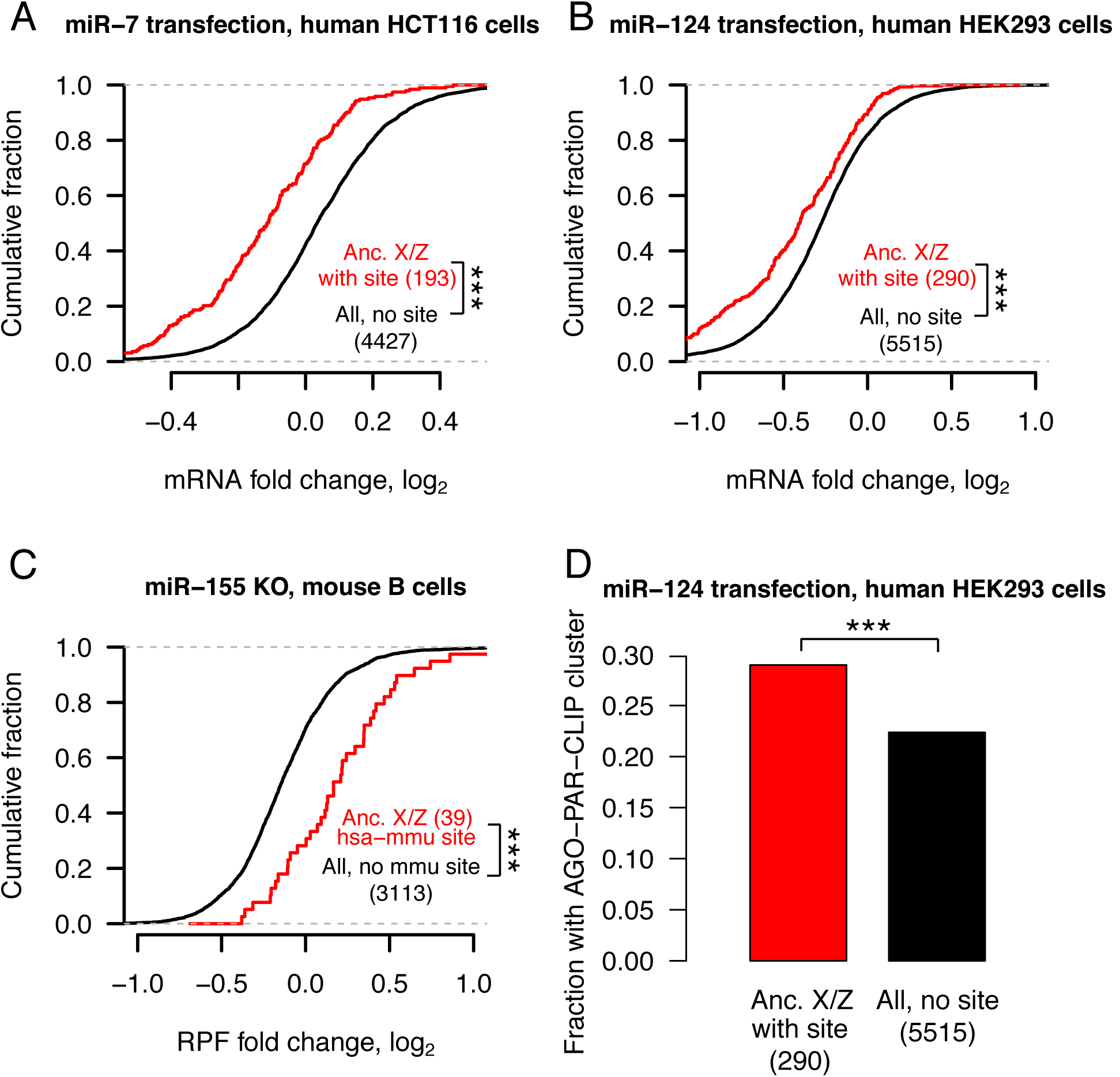
Analyses of experimental datasets validate miRNA target site efficacy. Distribution of changes in gene expression (A,B), in mRNA stability and translational efficiency as measured by ribosome protected fragments (RPF, C), and in fraction of Argonaute-bound genes (D) following miRNA transfection or knockout in the indicated cell-type. In each case, X-linked genes and the human orthologs of Z-linked genes containing target sites for the indicated miRNA were compared to all expressed genes lacking target sites; gene numbers are indicated in parentheses. (A-C) *** p < 0.001, two-sided Kolmogorov-Smirnov test. (D) *** p < 0.001, two-sided Fisher’s exact test.

## DISCUSSION

Here, through phylogenetic and functional analysis of conserved microRNA (miRNA) targeting, we demonstrate preexisting heterogeneities in dosage sensitivity among genes on the mammalian X and avian Z chromosomes. We first showed that, across all human autosomal genes, dosage sensitivity — as indicated by patterns of genic copy number variation — correlates with the degree of conserved miRNA targeting. Turning to the sex chromosomes of mammals and birds, genes that retained a homolog on the sex-specific Y or W chromosome (X-Y and Z-W pairs) are more dosage-sensitive than genes with no Y or W homolog. In mammals, genes with no Y homolog that became subject to XCI are more dosage-sensitive than those that continued to escape XCI following Y gene decay. These differences in dosage sensitivity were present on the ancestral autosomes that gave rise to the mammalian and avian sex chromosomes. Finally, through analysis of publically available experimental datasets, we validated the efficacy, in living cells, of the miRNA target sites used to infer dosage sensitivity. We thus conclude that differences in dosage sensitivity among genes on the ancestral autosomes influenced their evolutionary trajectory during sex chromosome evolution, not only on the sex-specific Y and W chromosomes, but also on the sex-shared X chromosome.

We and others previously proposed that Y gene decay drove upregulation of homologous X-linked genes in both males and females, and that XCI subsequently evolved at genes sensitive to increased expression from two active X-linked copies in females (Jegalian & Page, 1998; Ohno, 1967). Our finding that X-inactivated genes have higher miRNA conservation scores than X escape genes is consistent with this model. However, our finding of heterogeneities in dosage sensitivity among the ancestral, autosomal precursors of both mammalian X-linked and avian Z-linked genes challenges the previous assumption of a single evolutionary pathway for all sex-linked genes.

We therefore propose a model of X-Y and Z-W evolution in which the ancestral autosomes that gave rise to the mammalian and avian sex chromosomes contained three (or two, in the case of birds) classes of genes with differing dosage sensitivities (Figure 7A,B). For ancestral genes with high dosage sensitivity, Y or W gene decay would have been highly deleterious, and thus the Y- or W-linked genes were retained. According to our model, these genes’ high dosage sensitivity also precluded upregulation of the X- or Z-linked homolog, and, in mammals, subsequent X-inactivation; indeed, their X-linked homologs continue to escape XCI (Bellott et al., 2014). For ancestral mammalian genes of intermediate dosage sensitivity, Y gene decay did occur, and was accompanied or followed by compensatory upregulation of the X-linked homolog in both sexes; the resultant increased expression in females was deleterious and led to the acquisition of XCI. Ancestral mammalian genes of low dosage sensitivity continued to escape XCI following Y decay, and perhaps did not undergo compensatory X upregulation (Figure 6A). These genes’ dosage insensitivity set them apart biologically, and evolutionarily, from the other class of X-linked genes escaping XCI — those with a surviving Y homolog.

**Figure 7.**
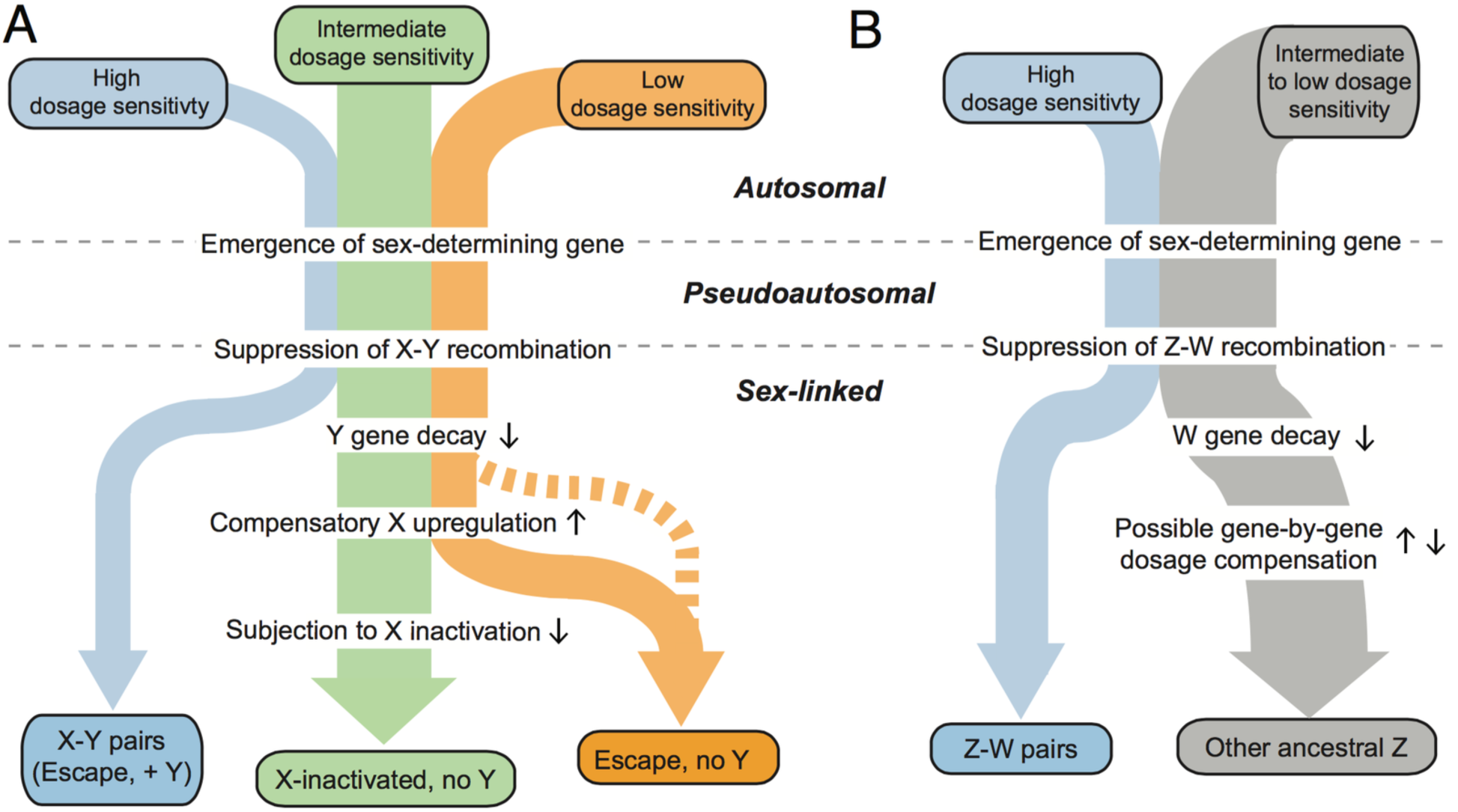
A proposed model of mammalian and avian sex chromosome evolution from ancestral autosomes. In this model, preexisting heterogeneities in dosage sensitivity determined the trajectory of Y/W gene loss in both mammals and birds and of subsequent X-inactivation in mammals. Colored arrow widths are scaled approximately to the number of ancestral genes in each class. The dashed orange line represents the possibility that a subset of X-linked genes may have not undergone compensatory X upregulation following Y gene decay. Small black arrows adjacent to molecular evolutionary transitions indicate increases or decreases in sex-linked gene dosage.

Previous studies have sought evidence of X-linked upregulation, during mammalian sex chromosome evolution, through comparisons of gene expression levels between the whole of the X chromosome and all of the autosomes, with equal numbers of studies rejecting or finding evidence consistent with upregulation (Deng et al., 2011; Julien et al., 2012; Kharchenko, Xi, & Park, 2011; Lin, Xing, Zhang, & He, 2012; Xiong et al., 2010). We find these difficulties unsurprising given the gene-by-gene heterogeneity in dosage sensitivities revealed by our present study. As mentioned above, X upregulation may have not evolved for genes of low dosage sensitivity. Among genes that did evolve X upregulation, gene-by-gene variation in the evolutionary timing of the upregulation – determined by the time of Y gene decay and the relative dosage sensitivity of the X-linked homolog – could result in more recently upregulated genes showing a stronger signature of upregulation as assayed by current expression levels. Furthermore, constraint on X-linked gene dosage may vary across tissues, such that X upregulation evolved in different tissues for different genes. Such variation across both genes and tissues could confound approaches that assume either the presence or absence of a uniform, chromosome-wide signature of X upregulation within individual tissues. In avian lineages, heterogeneity in dosage sensitivities among the autosomal precursors of Z-linked genes, which our present study shows to be significant factor in W gene retention, may have caused Z dosage compensation to evolve on a gene-by-gene basis. Indeed, studies of Z-linked gene expression provide evidence for the gene-by-gene nature of Z dosage compensation (Itoh et al., 2007; Mank & Ellegren, 2009; Uebbing et al., 2015). In contrast to studies focusing on the readily apparent differences in dosage compensation mechanism between mammals and birds, our mechanistic findings and accompanying model highlight an aspect of dosage compensation of sex-linked genes that encompasses both birds and mammals.

In addition to highlighting similarities between mammals and birds, our study provides a view of dosage compensation that highlights post-transcriptional regulatory mechanisms. Previous studies have drawn inferences regarding post-transcriptional dosage compensation from measurements of extant protein abundance (Chen & Zhang, 2015; Uebbing et al., 2015). Our study points to specific non-coding sequences with known mechanisms (microRNA target sites) functioning across evolutionary time. Perhaps additional post-transcriptional regulatory mechanisms and their associated regulatory elements will be revealed to play roles in mammalian and avian dosage compensation.

Human disease studies support our model’s claim that increased dosage of X-Y pairs and X-inactivated genes is deleterious to fitness. Copy number gains of the X-linked gene *KDM6A*, which has a surviving human Y homolog, are found in patients with developmental abnormalities and intellectual disability (Lindgren et al., 2013). *HDAC6*, *CACNA1F*, *GDI1*, and *IRS4* all lack Y homologs and are subject to XCI in humans. A mutation in the 3’ UTR of *HDAC6* abolishing targeting by miR-433 has been linked to familial chondrodysplasia in both sexes (Simon et al., 2010). Likely gain-of-function mutations in *CACNA1F* cause congenital stationary night blindness in both sexes (Hemara-Wahanui et al., 2005). Copy number changes of *GDI1* correlate with the severity of X-linked mental retardation in males, with female carriers preferentially inactivating the mutant allele (Vandewalle et al., 2009). Somatic genomic deletions downstream of *IRS4* lead to its overexpression in lung squamous carcinoma (Weischenfeldt et al., 2017). Males with partial X disomy due to translocation of the distal long arm of the X chromosome (Xq28) to the long arm of the Y chromosome show severe mental retardation and developmental defects (Lahn et al., 1994). Most genes in Xq28 are inactivated in 46,XX females but escape inactivation in such X;Y translocations, suggesting that increased dosage of Xq28 genes caused the cognitive and developmental defects. We anticipate that further studies will reveal additional examples of the deleterious effects of overexpression of X-Y pairs and X-inactivated genes.

Recent work has revealed that the sex-specific chromosome — the Y in mammals and the W in birds — convergently retained dosage-sensitive genes with important regulatory functions (Bellott et al., 2014, 2017). Our study provides direct evidence that such heterogeneity in dosage sensitivity existed on the ancestral autosomes that gave rise to the mammalian and avian sex chromosomes. This heterogeneity influenced both survival on the sex-specific chromosomes in mammals and birds and the evolution of XCI in mammals. Thus, two independent experiments of nature have shown that modern-day amniote sex chromosomes were shaped, during evolution, by the properties of the ancestral autosomes from which they derive.

## METHODS

### Human genic copy number variation

To annotate gene deletions and duplications, we used data from the Exome Aggregation Consortium (ExAC, RRID:SCR_004068)

(ftp://ftp.broadinstitute.org/pub/ExAC_release/release0.3.1/cnv/), which consists of autosomal genic duplications and deletions (both full and partial) called in 59,898 exomes (Ruderfer et al., 2016). We used the publicly available genic deletion counts but re-computed genic duplication counts using only full duplications, reasoning that partial duplications are unlikely to result in increased dosage of the full gene product. We thus required that an individual duplication fully overlapped the longest protein-coding transcript (GENCODE v19) of a gene using BEDtools (RRID:SCR_006646) (Quinlan & Hall, 2010). We removed genes flagged by ExAC as lying in known regions of recurrent CNVs. This yielded 4,118 genes within duplications and deletions, 3,976 genes within deletions but not duplications, 2,916 genes within duplications but not deletions, and 3,510 genes not subject to duplication or deletion.

### X-linked gene sets

Analyses of conserved miRNA targeting based on multiple species alignments are unreliable for multicopy or ampliconic genes due to ambiguous sequence alignment between species. To avoid such issues, we first removed multicopy and ampliconic genes (Mueller et al., 2013) from a previously published set of human X genes present in the amniote ancestor (Bellott et al., 2014). We then excluded genes in the human pseudoautosomal (PAR) regions since these genes have not been exposed to the same evolutionary forces as genes in regions where X-Y recombination has been suppressed. Of the remaining ancestral X genes, we classified the 15 genes with human Y-linked homologs as X-Y pairs. We also analyzed the larger set of 32 X-Y pairs across eight mammals (human, chimpanzee, rhesus macaque, marmoset, mouse, rat, bull, and opossum) with sequenced Y chromosomes (Bellott et al., 2014).

To classify ancestral X-linked genes without Y homologs as subject to or escaping XCI in humans, we used a collection of consensus XCI calls which aggregate the results of three studies (Carrel & Willard, 2005; Cotton et al., 2013, 2015) assaying XCI escape (Balaton et al., 2015). Out of 472 ancestral X genes without a human Y homolog assigned an XCI status by Balaton et al. (Balaton et al., 2015), 329 were subject to XCI (“Subject” or “Mostly subject” in Balaton et al.), 26 displayed variable escape (“Variable escape” or “Mostly variable escape”) from XCI, and 30 showed consistent escape (“Escape” or “Mostly escape”). We excluded 40 ancestral X genes with a “Discordant” XCI status as assigned by Balaton et al. In the main text, we present results obtained after combining both variable and consistent escape calls from Balaton et al. into one class, yielding the following counts: 15 X-Y pairs, 329 ancestral X genes subject to XCI, and 56 ancestral X genes with evidence of escape from XCI. We also performed analyses considering escape and variable escape genes separately (Figure 2 – figure supplement 3).

### Z-linked gene sets

We previously refined the ancestral gene content of the avian sex chromosomes to 685 Z-linked genes with human orthologs by sequencing of the chicken Z chromosome and analysis of 13 other avian species with published female genomes (Bellott et al., 2017). Of these 685 ancestral Z genes, 28 retained a homolog on the fully sequenced chicken W chromosome. Including three additional avian species in which candidate W-linked genes were ascertained by directly comparing male and female genome assemblies results in a total of 78 W-linked genes. Including another 10 avian species in which W-linkage was inferred by read depth changes in a female genome results in a total of 157 W-linked genes.

### microRNA target site P_CT_ scores

Summaries of all gene-miRNA family interactions were obtained from TargetScanHuman v7.1 (RRID:SCR_010845)(http://www.targetscan.org/vert_71/vert_71_data_download/Summary_Counts.all_predictions.txt.zip). We excluded mammalian-specific miRNA families based on classifications by Friedman et al (Friedman et al., 2009)and updated in TargetScanHuman v7.1(Agarwal, Bell, Nam, & Bartel, 2015). To account for gene-specific variability in the number and P_CT_ score of gene-miRNA interactions within a group of genes, we sampled 1000x with replacement from the same group of genes and computed the mean gene-miRNA P_CT_ score for all associated gene miRNA interactions from each sampling. These 1000 samplings were then used to estimate the median resampled gene-miRNA P_CT_ and 95% confidence intervals. To compare P_CT_ scores between two sets of genes while matching by haploinsufficiency probability (Huang et al., 2010), we binned the smaller of the two gene sets into quintiles based on the assigned haploinsufficiency probabilities. We then binned the second gene set based on these same quintiles. For each of 1000 samplings, we randomly drew *n* genes from the second set from each of the quintile bins, where *n* is the number of genes from the first set assigned to that quintile. We computed a mean gene-miRNA P_CT_ score from each randomly sampled gene set and used this distribution to compute an empirical one-sided P-value.

### Human-chicken conserved microRNA target sites

Site-wise alignment information was obtained from TargetScanHuman v7.1 (http://www.targetscan.org/vert_71/vert_71_data_download/Conserved_Family_Info.txt.zip). To determine which target sites are present in the 3’ UTRs of both human and chicken orthologs, we counted, for genes with both a human and chicken ortholog, the number of miRNA interactions that had at least one target site in both human and chicken. To control for gene-specific background 3’ UTR conservation, we generated six control k-mers for each miRNA family seed sequence that were matched exactly for nucleotide and CpG content. Six was the maximum number of unique control k-mers that could be generated for all sequences. We repeated the above counting analysis with each of the control k-mers using scripts from TargetScan, and compared, for each gene, the observed number of human-chicken-conserved miRNA interactions (the observed conservation signal) to the average number from controls (the background conservation).

### Gene expression profiling and crosslinking datasets

Changes in mRNA expression from a compendium of small RNA (sRNA) transfections (corresponding to twelve different miRNAs) in HeLa cells were obtained from Agarwal and colleagues, who carefully normalized microarray data (Agarwal et al., 2015). Further datasets describing the effects of transfecting miR-103 in HCT116 cells (Linsley et al., 2007), knocking down miR-92a in HEK293 cells (Hafner et al., 2010), transfecting miR-7 or miR-124 in HEK293 cells (Hausser, Landthaler, Jaskiewicz, Gaidatzis, & Zavolan, 2009), or of knocking out miR-155 in mouse B cells (Eichhorn et al., 2014), T cells (Loeb et al., 2012), or Th1 and Th2 cells (Rodriguez et al., 2007), processed as described in (Agarwal et al., 2015), were provided by

V. Agarwal. Targets for the PAR-CLIP study (Hafner et al., 2010) were inferred from an online resource of HEK293 clusters observed after transfection of either miR-124 (http://www.mirz.unibas.ch/restricted/clipdata/RESULTS/miR124_TRANSFECTION/miR124_TRANSFECTION.html) or miR-7 (http://www.mirz.unibas.ch/restricted/clipdata/RESULTS/miR7_TRANSFECTION/miR7_TRANSFECTION.html).

## Data availability

Data supporting the findings of this study are available within the paper and its supplementary information files.

## Code availability

Custom scripts in R (RRID:SCR_001905) were used to generate figures. A custom Python (RRID:SCR_008394) script utilizing Biopython (RRID:SCR_007173) was used to generate shuffled miRNA family seed sequences; all code is available upon request from the authors. Identification of miRNA target site matches using shuffled seed sequences was performed using the ‘targetscan_70.pl’ perl script (http://www.targetscan.org/vert_71/vert_71_data_download/targetscan_70.zip).

## Author contributions

S.N., D.W.B. and D.C.P designed the study. S.N. performed analyses with assistance from D.W.B. S.N. and D.C.P wrote the paper.

## Acknowledgements

We thank V. Agarwal, S. Eichorn, S. McGeary, and D. Bartel for assistance with the TargetScan database and helpful discussions; A. Godfrey for updated human-chicken orthology information; and A. Godfrey, J. Hughes and H. Skaletsky for critical reading of the manuscript. This work was supported by the National Institutes of Health and the Howard Hughes Medical Institute. S.N. was supported under a research grant by Biogen.

---

**Figure 1– Figure Supplement 1:**
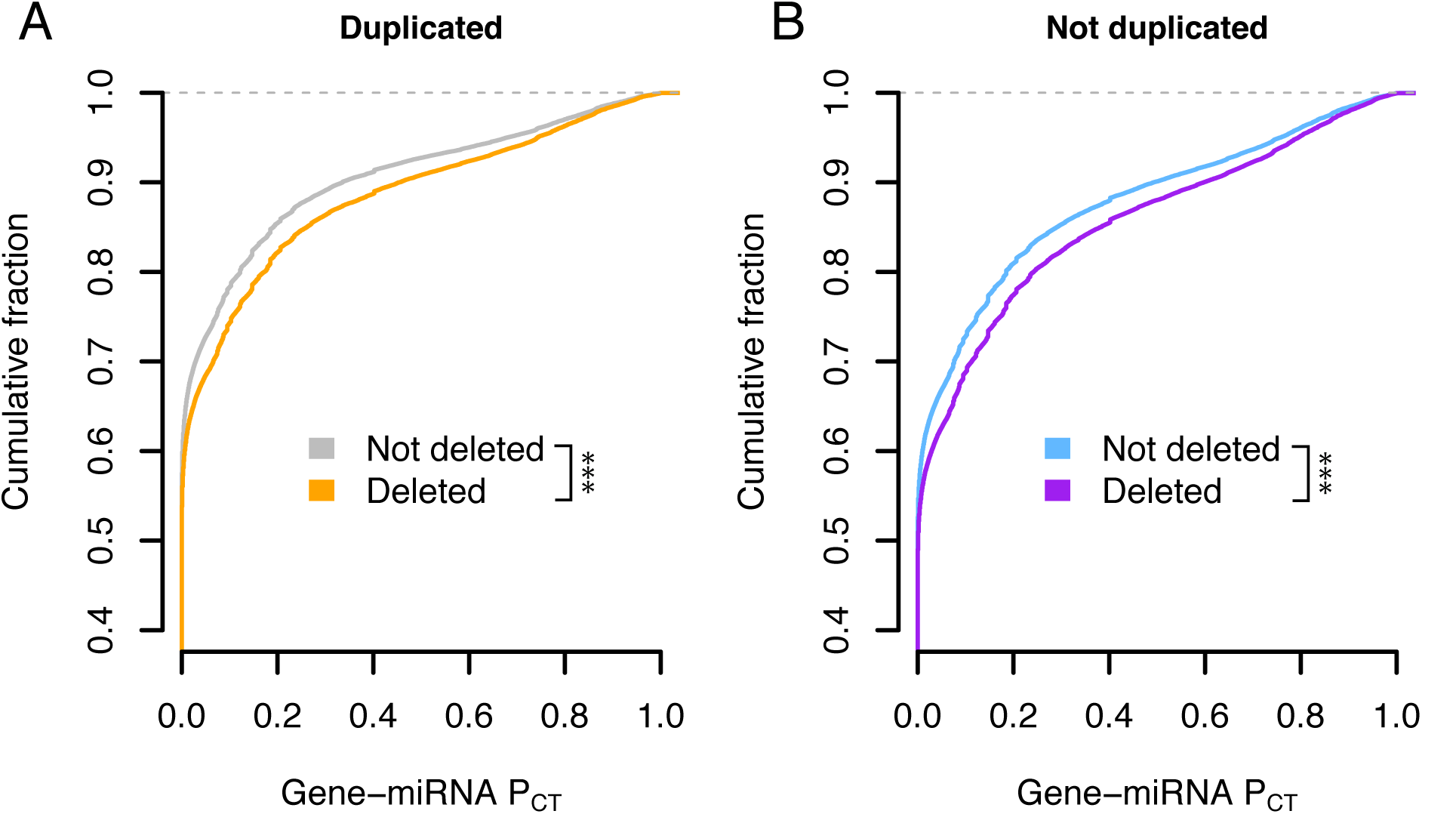
Effect of deletion status on autosomal P_CT_ scores. Probabilities of conserved targeting (PCT) of all gene-miRNA interactions involving non-deleted and deleted genes, further stratified as (A) duplicated (grey, n = 69,339 interactions from 4,118 genes; orange, n = 51,514 interactions from 2,916 genes) or (B) not duplicated (purple, n = 72,826 interactions from 3,510 genes; blue, n = 80,290 interactions from 3,976 genes). *** p < 0.001, two-sided Kolmogorov-Smirnov test.

**Figure 1 – Figure Supplement 2:**
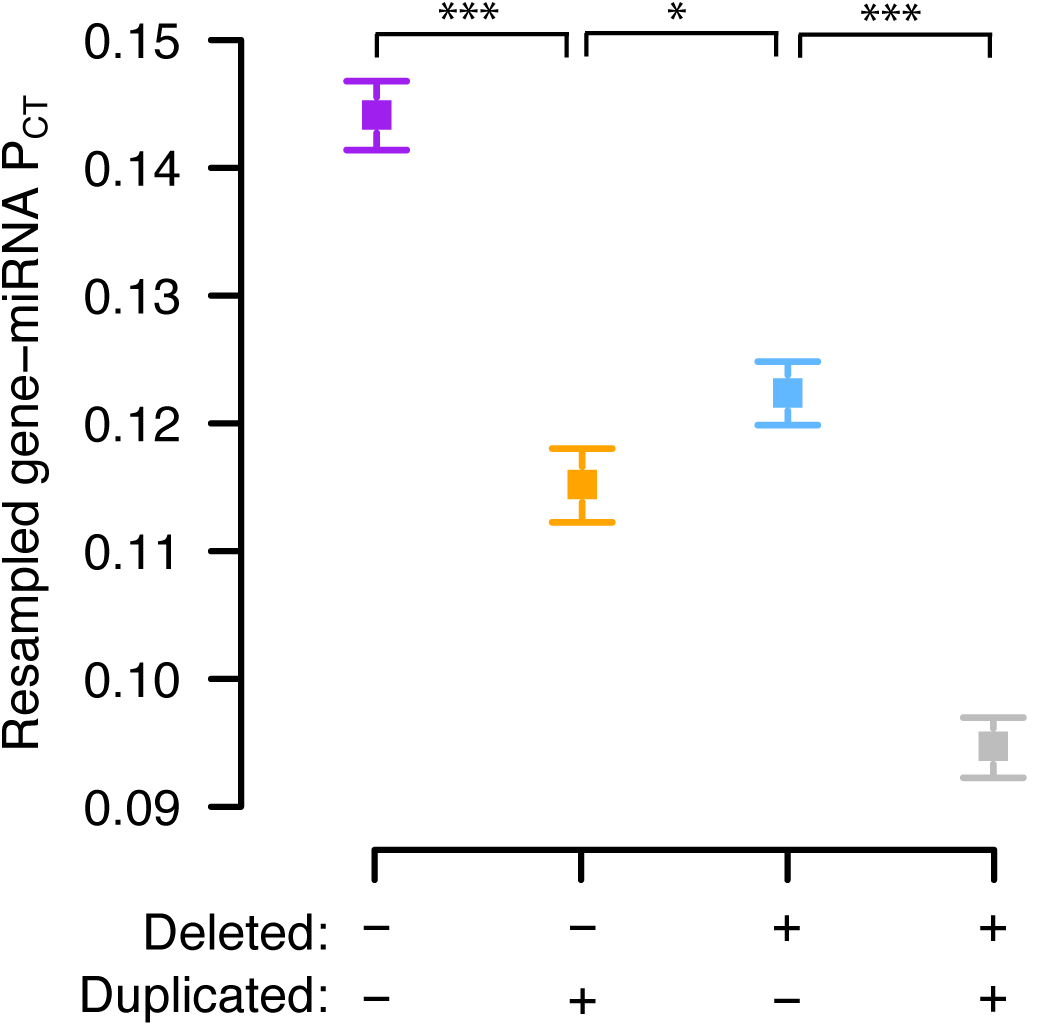
Resampled mean P_CT_ scores of autosomal genes stratified by copy number variation in 59,898 human exomes. Non-duplicated and duplicated genes were further stratified as deleted (blue, n = 3,976 genes; grey, n = 4,118 genes) or not deleted (purple, n = 3,510 genes; orange, n = 2,916 genes). Points and error bars represent the median and 95% confidence intervals of P_CT_ scores from 1,000 gene samplings with replacement. **\*** p < 0.05, *** p < 0.001, empirical p-value computed as the fraction of random non-overlapping gene sets with a median difference in P_CT_ score at least as large as the true difference.

**Figure 2–Figure Supplement 1:**
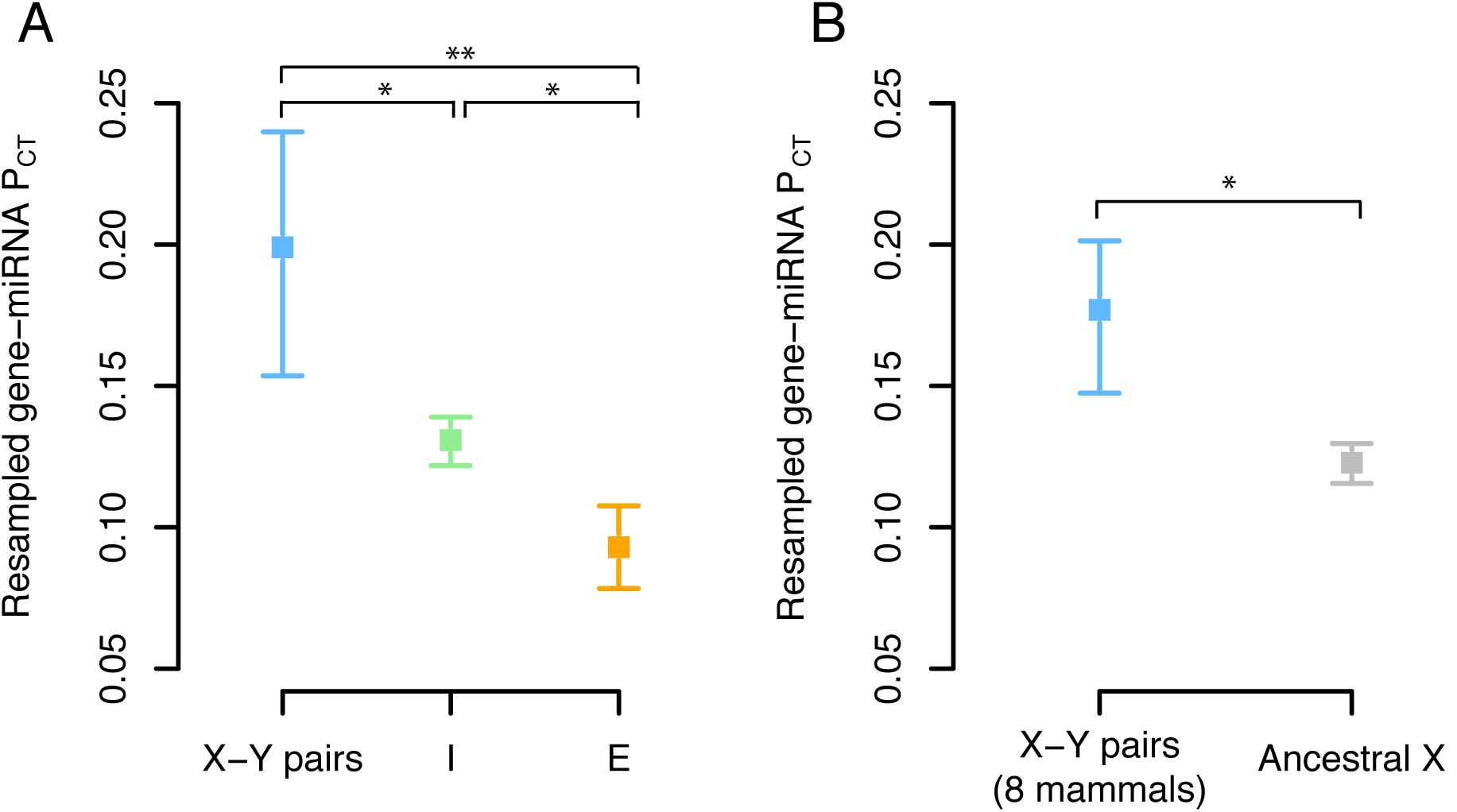
Resampled mean P_CT_ scores of X-linked genes. (A) Resampled gene-miRNA P_CT_ scores for human X-Y pairs (n = 15 genes), X-inactivated genes (n = 329 genes) and X escape genes (n = 56 genes). (B) Resampled gene-miRNA P_CT_ scores for X-Y pairs across eight mammals (n = 32 genes) and genes with no Y homolog in any of eight mammals (n = 457 genes). Points and error bars represent the median and 95% confidence intervals from 1,000 gene samplings with replacement. * p < 0.05, ** p < 0.01, empirical p-value computed as the fraction of random non-overlapping gene sets with a median difference in P_CT_ score at least as large as the true difference.

**Figure 2 – Figure Supplement 2:**
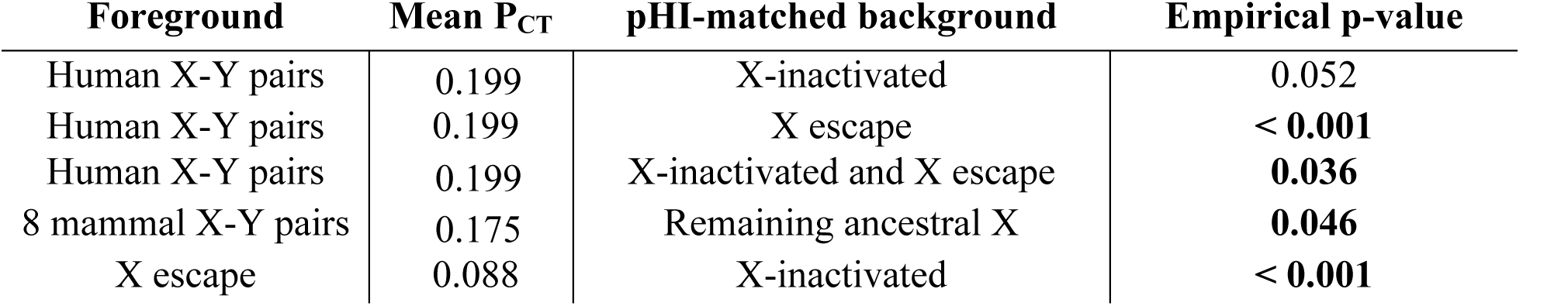
Resampling tests controlling for differences in haploinsufficiency probability between X-linked genes. In each row, genes from the “background” set (indicated in the third column) were matched to genes in the “foreground” set (first column) by their predicted haploinsufficiency probability. The mean P_CT_ score of the foreground set was compared to the distribution of mean P_CT_ scores from the matched background set to obtain a one-sided empirical p-value.

**Figure 2 – Figure Supplement 3:**
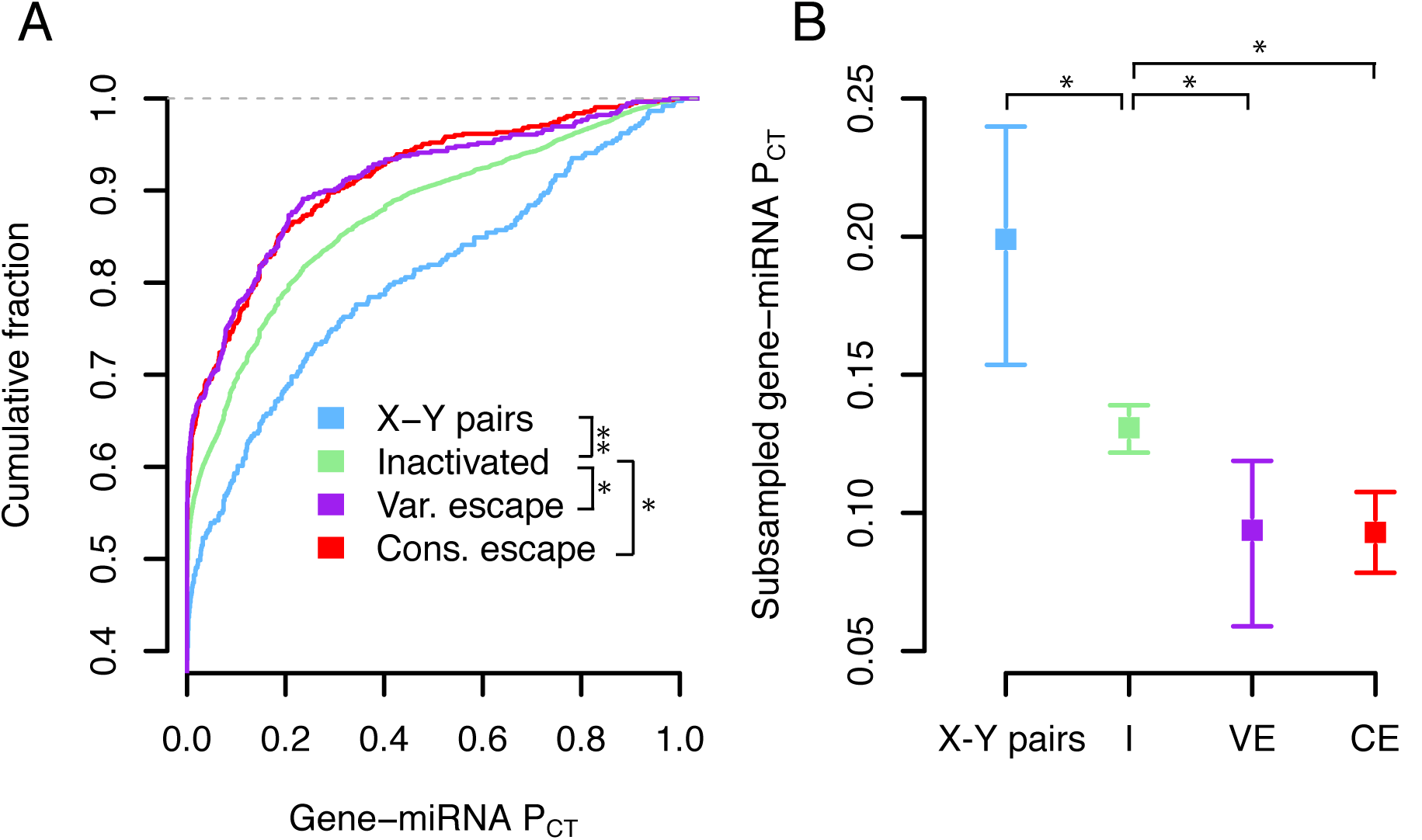
P_CT_ score comparisons with consistent and variable escape genes separated. (A) P_CT_ score distributions of all gene-miRNA interactions involving X-Y pairs (n = 371 interactions from 16 genes), X-inactivated genes (n = 6743 interactions from 329 genes), consistent escape genes (n = 567 interactions from 30 genes), or variable escape genes (n = 470 interactions from 26 genes) as defined by Balaton et al (Balaton et al., 2015). * p < 0.05, ** p < 0.01, two-sided Kolmogorov-Smirnov test. (B) Resampled gene-miRNA P_CT_ scores of gene classes from (A). Points and error bars represent the median and 95% confidence intervals from 1,000 gene samplings with replacement. * p < 0.05, empirical p-value computed as the fraction of random non-overlapping gene sets with a median difference in P_CT_ score at least as large as the true difference.

**Figure 3 – Figure Supplement 1:**
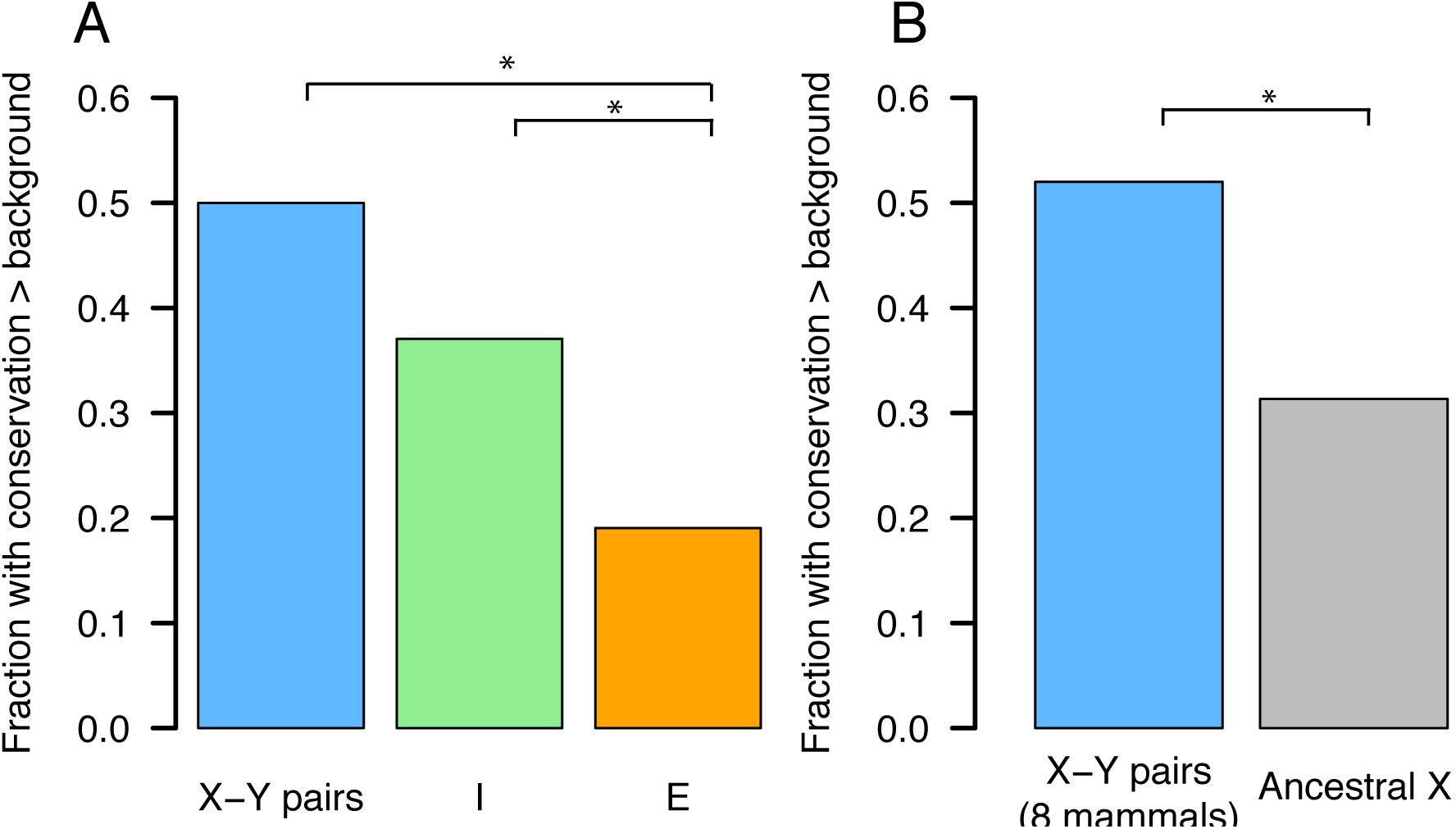
X-linked conservation of ancestral miRNA targeting above background levels. The fraction of genes in each class showing human-chicken miRNA target site conservation above background was calculated by comparing the true number of human-chicken-conserved target sites with the average number of human-chicken-conserved sites using six shuffled k-mers for each miRNA family. (A) Human X-Y pairs (n = 15 genes), X-inactivated genes (n = 329 genes) and X escape genes (n = 56 genes). (B) X-Y pairs across eight mammals (n = 32 genes), other ancestral X genes (n = 457 genes). * p < 0.05, two-sider Fisher exact test.

**Figure 4 – Figure Supplement 1:**
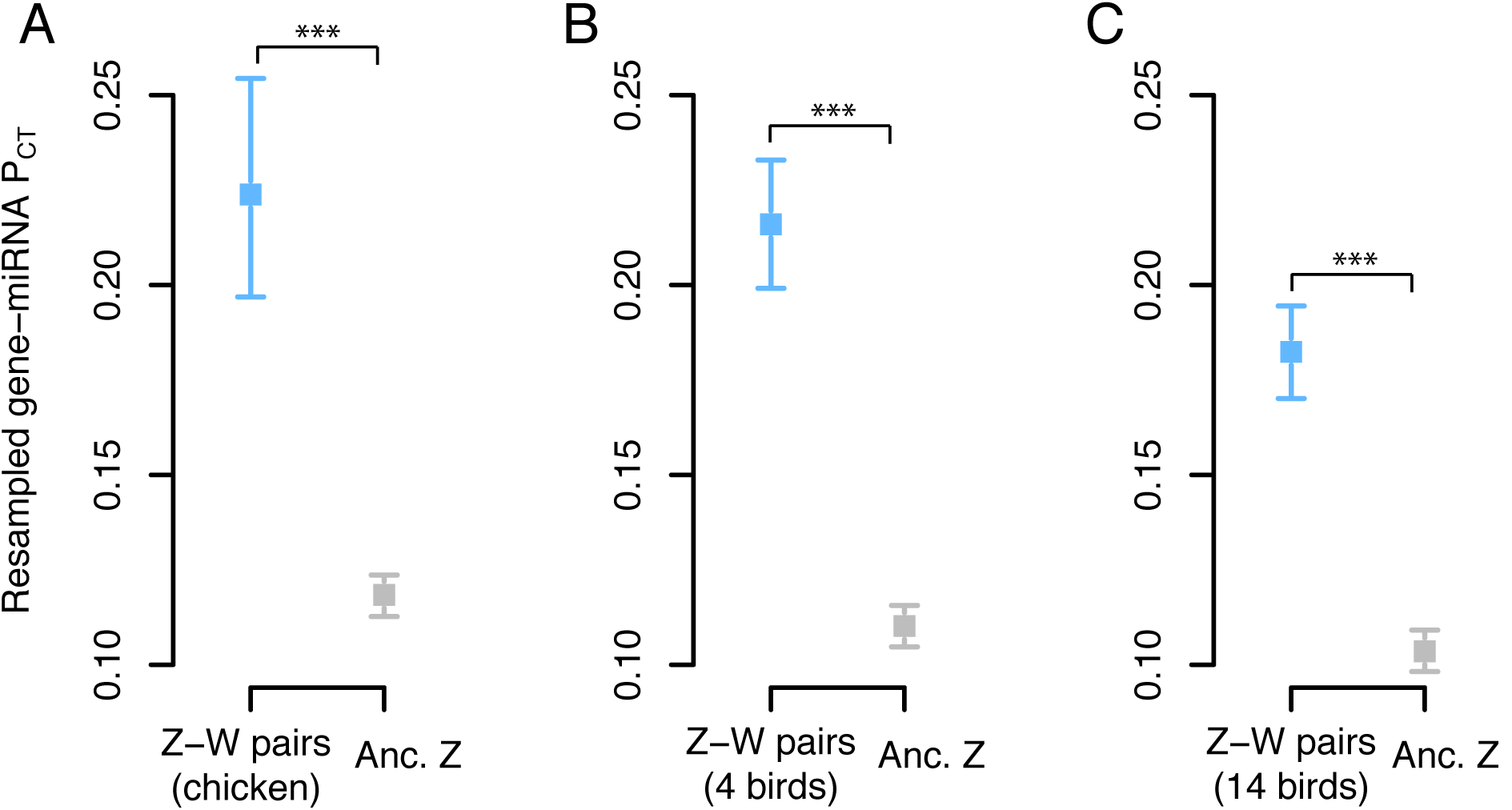
Resampled mean P_CT_ scores of Z-linked genes. Gene sets: (A) chicken Z-W pairs (n = 28 genes) and other ancestral Z genes (n = 657 genes), (B) Z-W pairs across four birds (n = 78 genes) compared to the remainder of ancestral Z genes (n = 607 genes), and (C) Z-W pairs across 14 birds (n = 157 genes) compared to the remainder of ancestral Z genes (n = 528 genes). Points and error bars represent the median and 95% confidence intervals from 1,000 gene samplings with replacement. *** p < 0.001, empirical p-value computed as the fraction of random non-overlapping gene sets with a median difference in PCT score at least as large as the true difference.

**Figure 4 – Figure Supplement 2:**
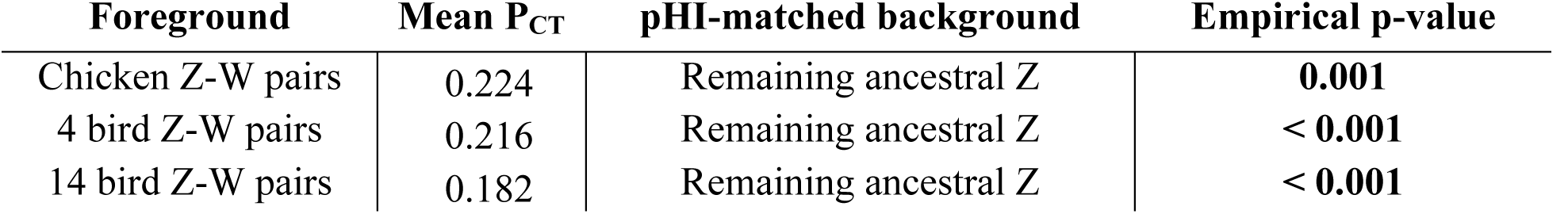
Resampling tests controlling for differences in haploinsufficiency probability between Z-linked genes. In each row, genes from the “background” set (indicated in the third column) were matched to genes in the “foreground” set (first column) by their predicted haploinsufficiency probability. The mean P_CT_ score of the foreground set was compared to the distribution of mean P_CT_ scores from the matched background set to obtain a one-sided empirical p-value.

**Figure 5 – Figure Supplement 1:**
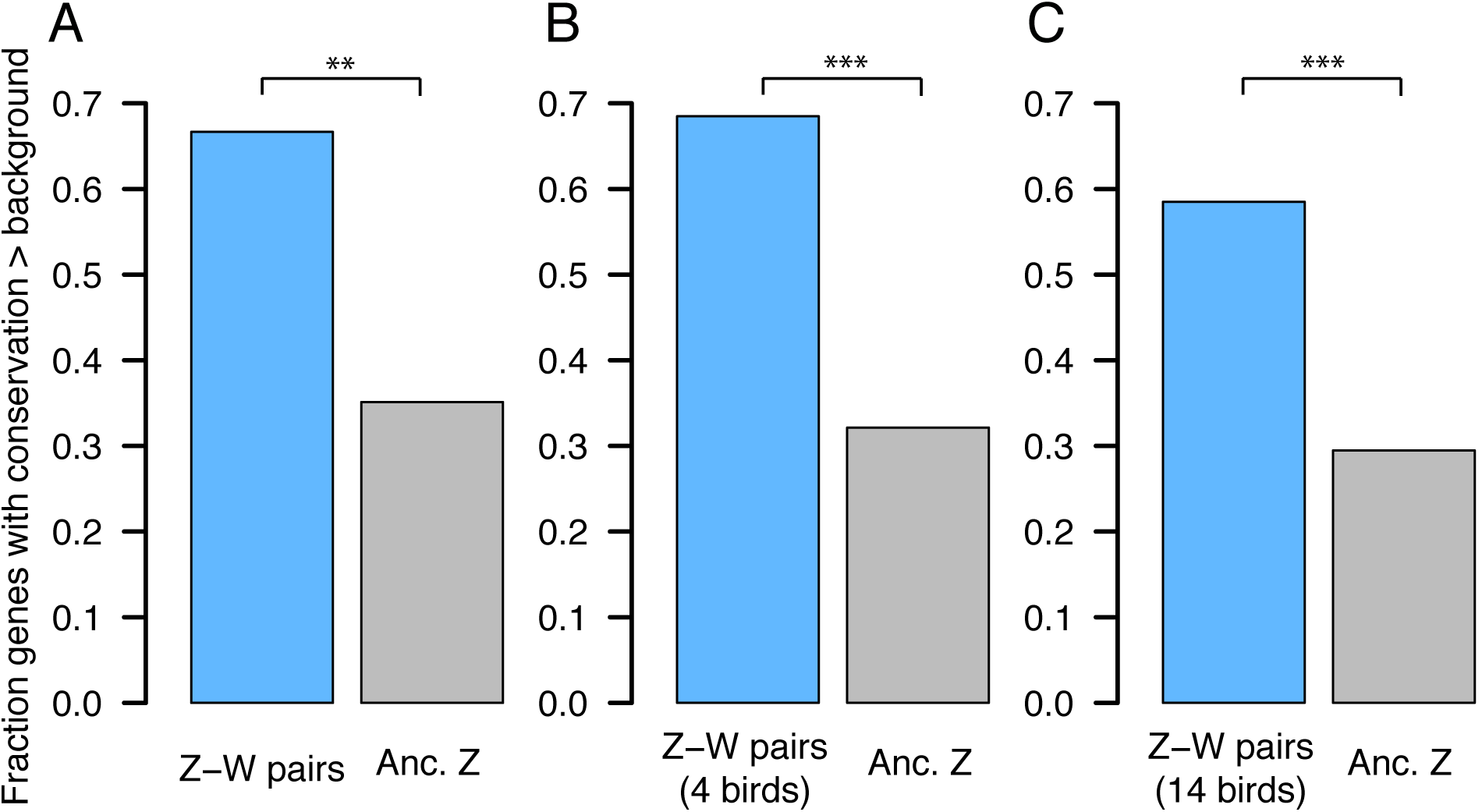
Z-linked conservation of ancestral miRNA targeting above background levels. The fraction of genes in each class showing human-chicken miRNA target site conservation above background was calculated by comparing the true number of human-chicken-conserved target sites with the average number of human-chicken-conserved sites using six shuffled k-mers for each miRNA family. (A) chicken Z-W pairs (n = 28 genes) and other ancestral Z genes (n = 657 genes), (B) Z-W pairs across four birds (n = 78 genes) compared to the remainder of ancestral Z genes (n = 607 genes), and (C) Z-W pairs across 14 birds (n = 157 genes) compared to the remainder of ancestral Z genes (n = 528 genes). ** p < 0.01, *** p < 0.001, two-sider Fisher exact test.

**Figure 6 – Figure Supplement 1:**
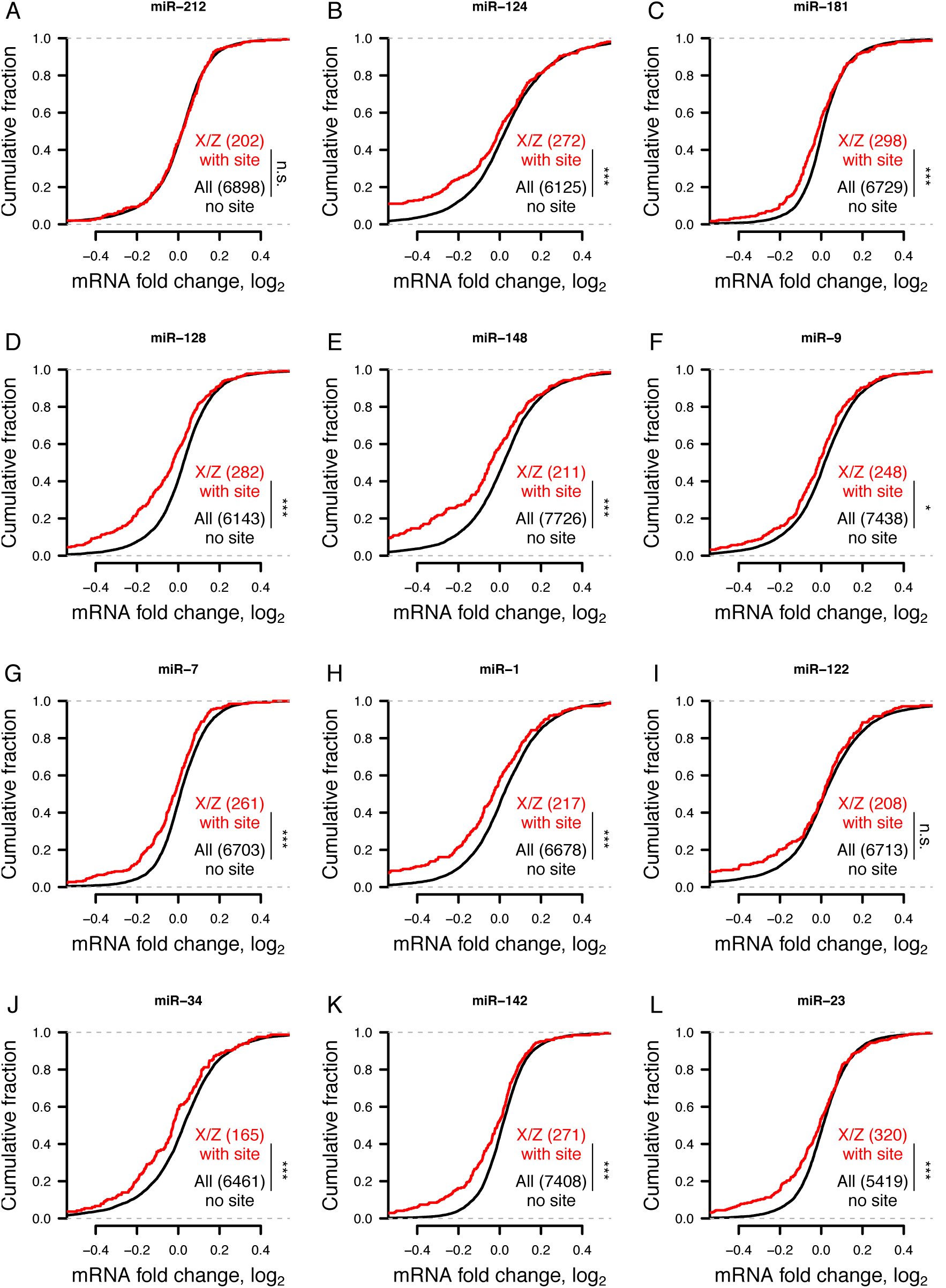
Gene expression changes following small RNA transfections in human HeLa cells. * p < 0.05, *** p < 0.001, two-sided K-S test.

**Figure 6 – Figure Supplement 2:**
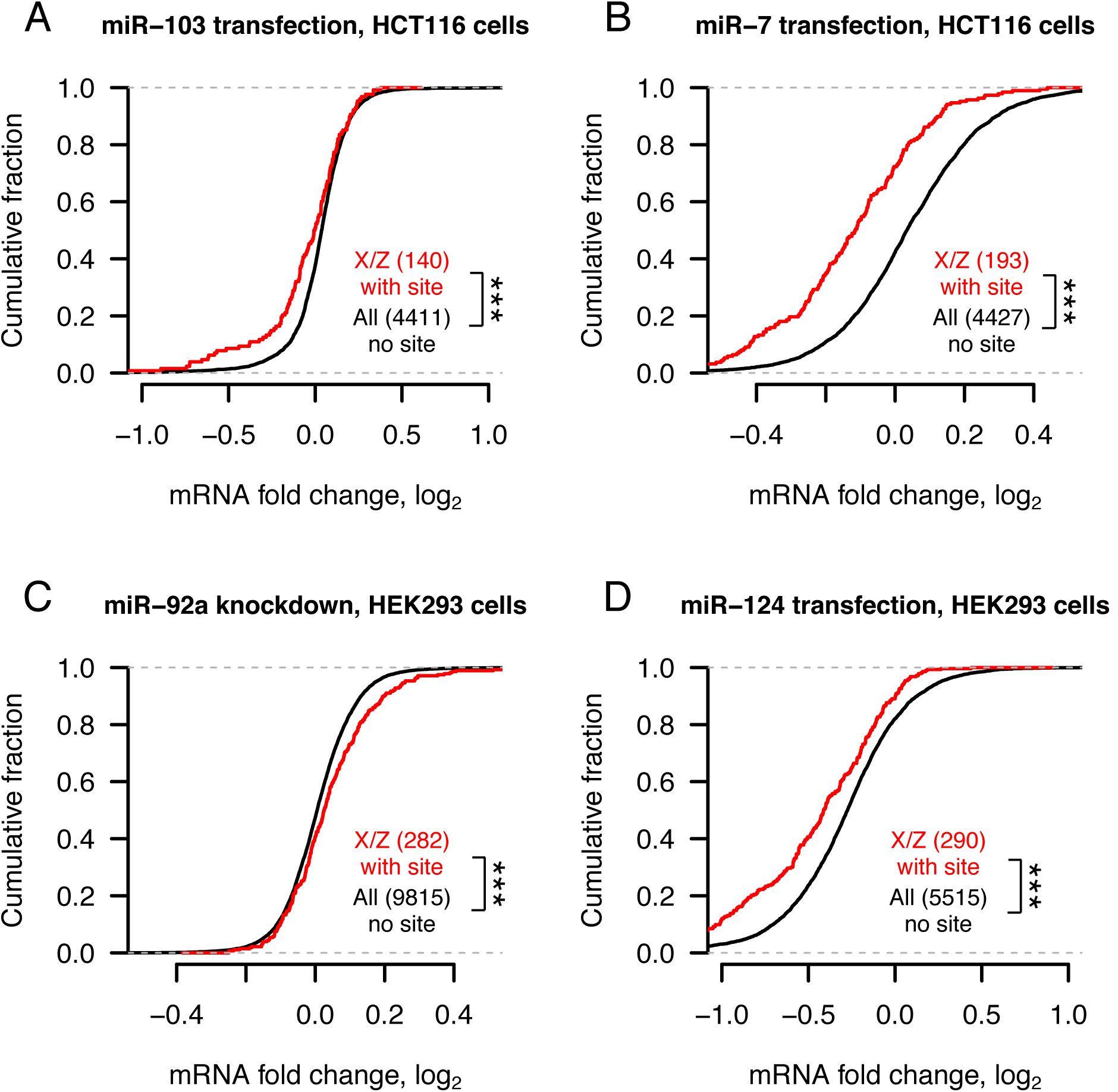
Gene expression changes following transfection or knockdown of additional miRNAs in human HCT116 or HEK293 cells. *** p < 0.001, two-side Kolmogorov-Smirnov test.

**Figure 6 – Figure Supplement 3:**
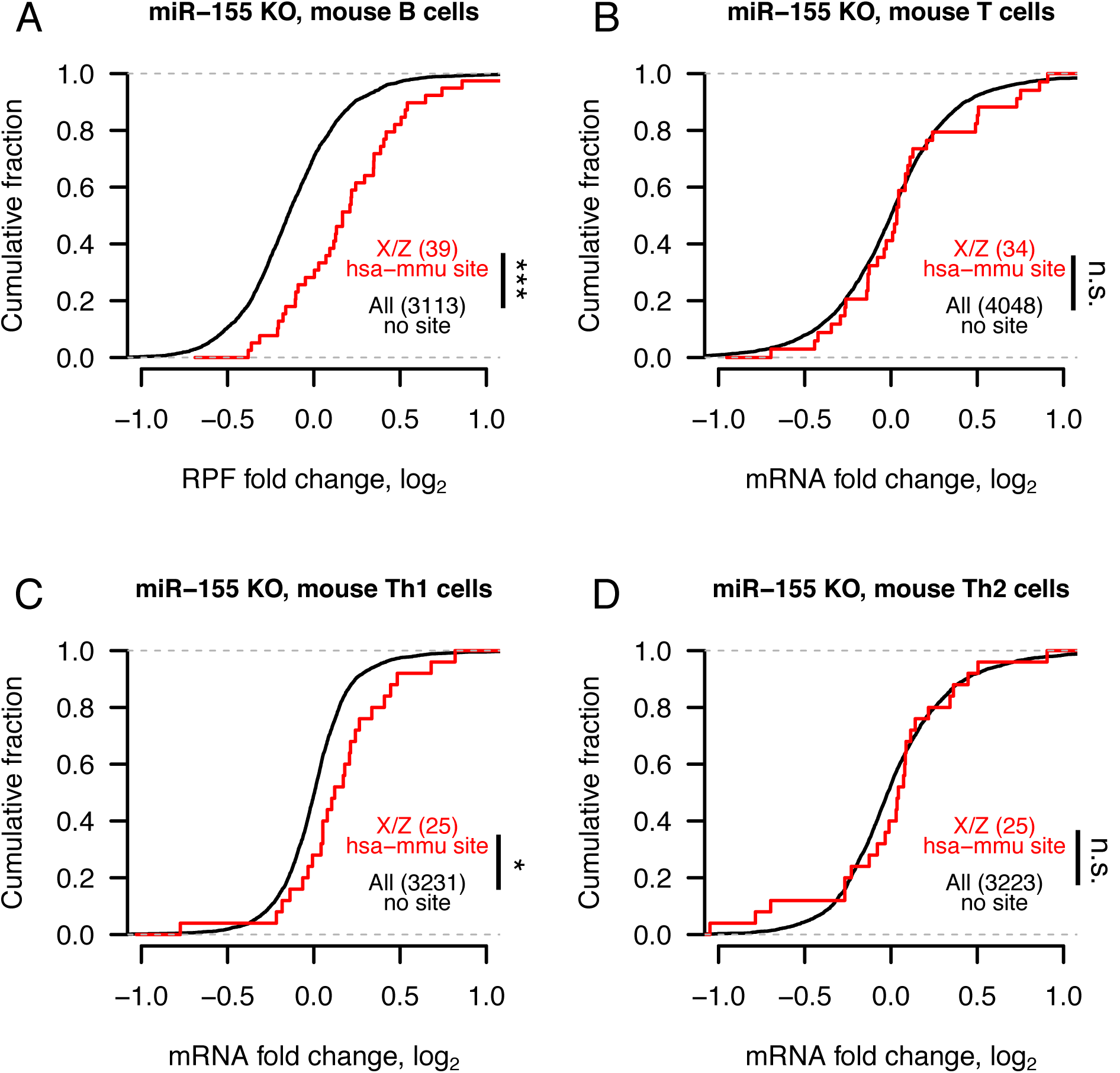
Changes in mRNA stability and translational efficiency and gene expression following miR-155 knockout in mouse immune cells. In each case, mouse orthologs of X- or Z-linked genes containing a human-mouse-conserved (hsa-mmu) miR-155 site were compared to human-mouse one-to-one orthologs lacking a miR-155 site. * p < 0.05, *** p < 0.001, two-sided Kolmogorov-Smirnov test.

**Figure 6 – Figure Supplement 4:**
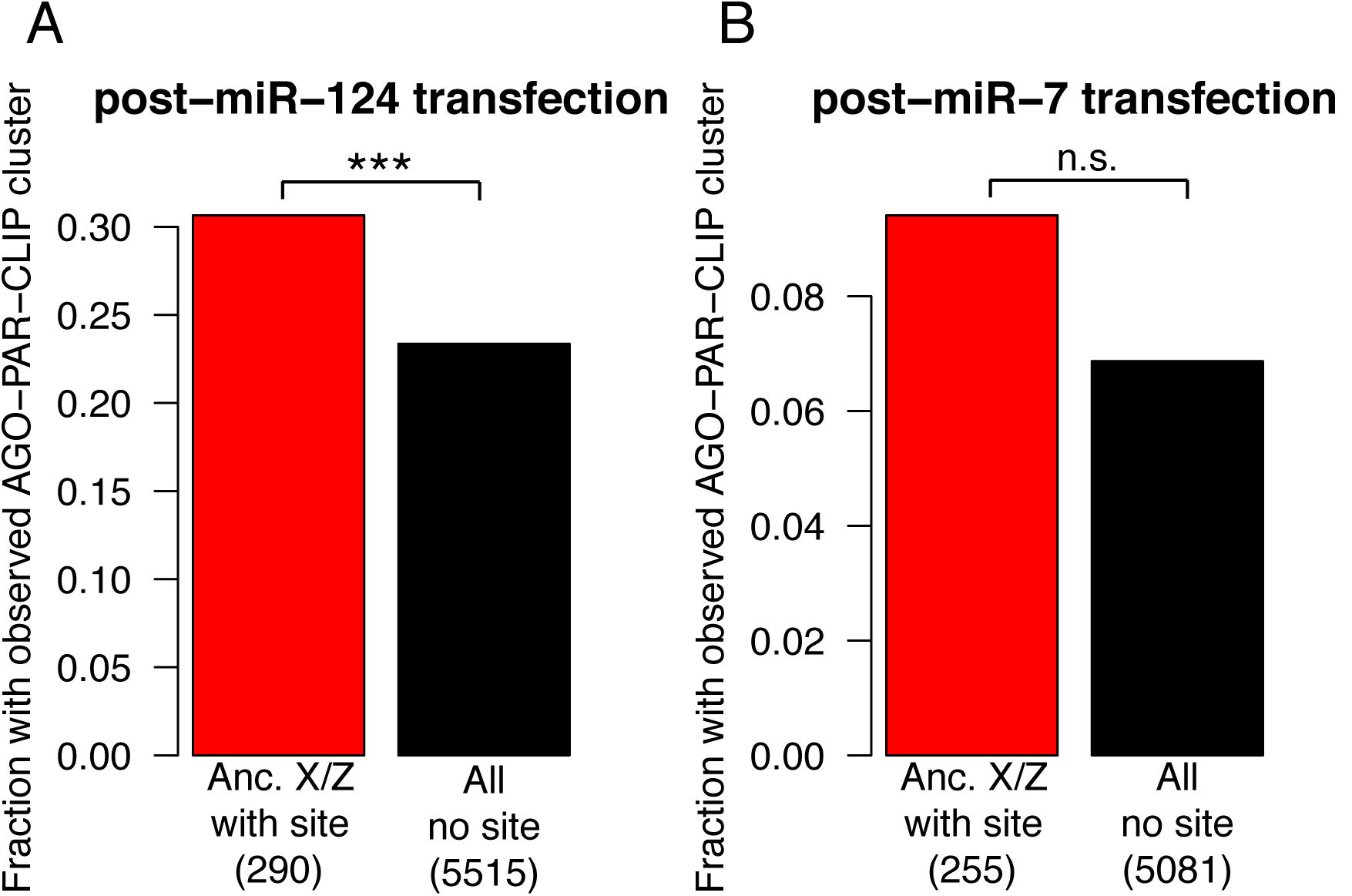
Argonaute binding measured by high-throughput crosslinking-immunoprecipitation (CLIP) following miRNA transfection in HEK293 cells. *** p < 0.001, two-sided Fisher’s exact test.

